# Optogenetic activators of apoptosis, necroptosis and pyroptosis for probing cell death dynamics and bystander cell responses

**DOI:** 10.1101/2021.08.31.458313

**Authors:** Kateryna Shkarina, Eva Hasel de Carvalho, José Carlos Santos, Maria Leptin, Petr Broz

## Abstract

Targeted and specific induction of cell death in individual or groups of cells holds the potential for new insights into the response of tissues or organisms to different forms of death. Here we report the development of optogenetically-controlled cell death effectors (optoCDEs), a novel class of optogenetic tools that enables light-mediated induction of three types of programmed cell death (PCD) – apoptosis, pyroptosis and necroptosis – using *Arabidopsis thaliana* photosensitive protein Cryptochrome2. OptoCDEs enable rapid and highly specific induction of PCD in human, mouse and zebrafish cells and are suitable for a wide range of applications, such as sub-lethal cell death induction or precise elimination of single cells or cell populations *in vitro* and *in vivo*. As the proof-of-concept, we utilize optoCDEs to assess the differences in the neighboring cell response to apoptotic or necrotic PCD, revealing a new role for shingosine-1-phosphate signaling in regulating the efferocytosis of apoptotic cell by epithelia.

## Introduction

Different types of programmed cell death (PCD) control the elimination of unnecessary, damaged, malignant or infected cells during development and tissue homeostasis ^1^. Apoptosis – the best studied form of PCD – leads to the disintegration of dying cells into apoptotic bodies that maintain membrane integrity and are removed by phagocytes in an immunologically silent manner^2^. Mechanistically, apoptosis is initiated either by the extrinsic/death-receptor pathway or the intrinsic/mitochondrial pathway that control the activation of apoptotic initiator and executioner caspases (caspase-8/-9, resp. caspase-3/-7), a family of cysteine-aspartic proteases that cleave substrate proteins to promote cell death. In contrast to apoptosis, programmed necrotic cell death such as pyroptosis and necroptosis result in cell lysis and the release of cytosolic content, which drives inflammation and immune responses^2^. Tumor necrosis factor receptor 1 (TNFR1) can initiate necroptosis if the activity of caspase-8 is inhibited by forming the necrosome, a signaling complex that contains the receptor-interacting kinase (RIP)-1 and RIP-3. Within this complex, RIP-1 phosphorylates RIP-3, which in turn phosphorylates and activates the pseudokinase MLKL that oligomerizes and translocates to the plasma membrane to form ion channels that drive necroptosis. Pyroptosis is initiated by inflammasome complexes^3^, large signaling hubs assembled upon detection of pathogens or endogenous danger signals by cytosolic pattern recognition receptors, resulting in activation of inflammatory caspases (e.g. caspase-1, −11, −4 and −5). These caspases process gasdermin D (GSDMD), thereby releasing the pore-forming GSDMD^NT^ that permeabilizes the plasma membrane to trigger pyroptosis.

Whereas the pathways and checkpoints that control PCD are well understood, comparably little is known about how the distinct forms of PCD differ in their outcome for the dying cell, its neighbors and the organism as a whole^4^. The effects of cell death are highly context specific and have been shown to either induce or dampen immune responses, or to induce cell proliferation and wound repair. In general, it is assumed that necrotic cell death causes inflammation, while apoptosis is anti-inflammatory or even tolerogenic^5^. Nevertheless, studies that systematically compare and contrast different forms of PCD and the respective response of neighboring cells remain rare, particularly in an *in vivo* setting. Experimentally, several obstacles hinder these analyses, such as the low specificity of cell death-inducing stimuli, which affect both the targeted cell as well as its neighbors, the crosstalk between different PCD pathways, and the difficulty to induce death in a spatially and temporally controlled manner. One strategy to overcome these obstacles is the use of laser-induced ablation of single or multiple cells^6–8^, as it allows to study the signaling response of direct neighbors and the extrusion of dead cells. However, it remains unclear what type of death is caused by laser ablation, since dying cells often display the features of both apoptosis (caspase activation) and necrosis (membrane permeabilization)^6^. Another more controlled approach is to force-oligomerize cell death executors using fusions with a chemically-dimerizable domain (DmrB/FKBP) (known as “clean” death system)^9^ but this is limited by poor reversibility and the inability to target and selectively kill individual cells *in vitro* or *in vivo*.

In this study, we present a novel set of ‘clean cell death’ tools for three major types of programmed cell death – pyroptosis, necroptosis and apoptosis – that is based on optogenetics (illumination with light); hence named <u>opto</u>genetically controlled <u>c</u>ell <u>d</u>eath <u>e</u>ffectors (optoCDEs). Compared to chemically inducible tools, light-mediated cell death induction offers several significant advantages: faster and easier signal delivery; ability to precisely control the intensity and duration of the cell death stimulus by varying the illumination dose and duration; and the ability to restrict cell death induction to selected cells or tissues of interest. These tools utilize Cryptochrome 2 E490G (Cry2olig), an *Arabidopsis thaliana*-derived photosensitive protein that undergoes rapid homooligomerization upon exposure to blue (450-488 nm) light^10^. By fusing Cry2olig to the effector domains of human, mouse or zebrafish inflammatory caspases, we designed light-activated caspases (optoCaspases), which induce rapid and efficient pyroptosis within minutes post illumination with blue light. We show that optoCaspases are functional in various model systems, including multiple human and mouse cell lines, organotypic 3D cell cultures and live zebrafish larvae, and can be utilized to precisely control, at a spatiotemporal level, the degree of caspase activation to drive sublytic pyroptosis or single-cell ablation. We further extend this approach to induce apoptosis and necroptosis, by generating light-activated apoptotic effectors optoCaspase-8 and optoCaspase-9 and necroptotic effectors optoRIP3 and optoMLKL. By using the optoCDEs, we demonstrate how optogenetic induction of cell death can be applied to study the response of bystander cells and the extrusion of dying cells from epithelia, revealing fundamental differences in the fates of cells dying by apoptosis and programmed necrosis.

## Results

### Design and characterization of optogenetically controlled human inflammatory caspases

In humans, pyroptosis is induced by the inflammatory caspases-1, −4 or −5 that cleave the pore-forming cell death executor GSDMD (**Fig. 1A**)^11^. Besides the catalytic p20 and p10 domains, inflammatory caspases feature a caspase recruitment domain (CARD), which mediates recruitment to inflammasome complexes and proximity-induced activation (**Fig. 1B**). To generate light-activatable caspases (hereafter termed optoCaspases), we fused the mCherry-tagged *A. thaliana* photosensitive protein Cry2olig N-terminally to either full-length or CARD-deficient human caspase-1, −4 or −5 (including different splice variants of caspase-5) (**Fig. 1C**) and transiently expressed these constructs in GSDMD-transgenic (GSDMD^tg^) HEK293T cells to assess their ability to induce pyroptosis upon blue light illumination. Light doses below 26.3 mW/cm^2^ (up to 3% of maximum 488 nm laser power) were used for all experiments where photoactivation was done by confocal microscopy, as this level of laser power did not cause detectable phototoxicity while efficiently inducing rapid Cry2olig clustering in HEK293T cells (**Supplementary Fig. 1A-B**). By contrast to Cry2olig expressing controls, optoCaspase-expressing (mCherry-positive) cells responded to blue light illumination by membrane blebbing, followed by cell swelling, membrane ballooning and nuclear rounding (**Fig. 1D-E, Supplementary Fig. 2A and Supplementary movie 1**), all of which are generally considered to be hallmarks of pyroptosis. These cells also rapidly lost membrane integrity, as measured by the influx of the otherwise membrane-impermeable DNA dye CellTox Green, acquired Annexin-V staining (measures phosphatidylserine (PS) exposure) and progressively lost cytoplasmic mCherry signal (**Fig. 1D and Supplementary Fig. 2B and Supplementary movie 1**). While all optoCaspase constructs induced death of more than 50% of mCherry^+^ cells within 30 min, full-length optoCaspase-1 and −5 were less efficient than the CARD-deficient variants. To better compare optoCaspase constructs and minimize potential endogenous regulatory interactions via the CARD, we decided to use CARD-deficient optoCaspase-1 (aa 92-404) and optoCaspase-4 (aa 92-377) and partially CARD-deficient optoCaspase-5 (aa 90-435) for all further experiments (of note, fully CARD-deficient optoCaspase-5 showed too high dark state cytotoxicity) (**Fig. 1D**). Kinetics of membrane permeabilization showed that these three constructs induced death with comparable speed and reached similar levels of pyroptosis at later timepoints, with the majority of optoCaspases-expressing cells (i.e. mCherry-positive) dying robustly within the first 10-15 min after the beginning of illumination (**Fig. 1E-F**). Mutation of the catalytic cysteines in optoCaspase-1, −4, or −5 completely abrogated cell death induction upon illumination (**Fig. 1G**), similarly to treatment with the pan-caspase inhibitor Z-VAD (**Fig. 1H**), validating the specificity of the tools. Mutating Asp387 of Cry2olig, which strongly decreases its light-mediated clustering^12^, also led to the reduction in pyroptosis levels (**Fig. 1G**). Light-induced activation of optoCaspase-1, −4 or −5 in wild-type HEK293T cells, which naturally lack endogenous GSDMD, did not induce membrane permeabilization and CellTox influx (**Supplementary Fig 2C-D**), since GSDMD is essential for pyroptosis induction by inflammatory caspases^11^. Instead, however, we observed the induction of a typical apoptotic morphology (**Supplementary Fig 2C-D**) that correlated with the activation of a genetically encoded caspase-3/7 reporter VC3AI^13^ (**Supplementary Fig. 2E**). This is in line with reports showing that caspase-1 activation in *Gsdmd*-deficient mouse macrophages engages rapid apoptosis via Bid cleavage^14–16^. Of note, while mouse caspase-11 is apparently not able to induce this type of apoptosis^7–9^, we found that optoCaspase-4 and −5 caused GSDMD-independent apoptosis, which could possibly be due to their wider substrate profile^17^. OptoCaspase activation efficiently induced pyroptosis in commonly used laboratory cell lines, such as MCF7 cells, HeLa, Caco-2, HT-29 and HaCaT, (**Fig. 1I and Supplementary Fig. 2F**). In summary, we demonstrate that fusion with the photoresponsive protein Cry2 can be used to drive inflammatory caspase activation and that optogenetically-activatable inflammatory caspases induce GSDMD-dependent pyroptosis across different cell types.

**Figure 1.**
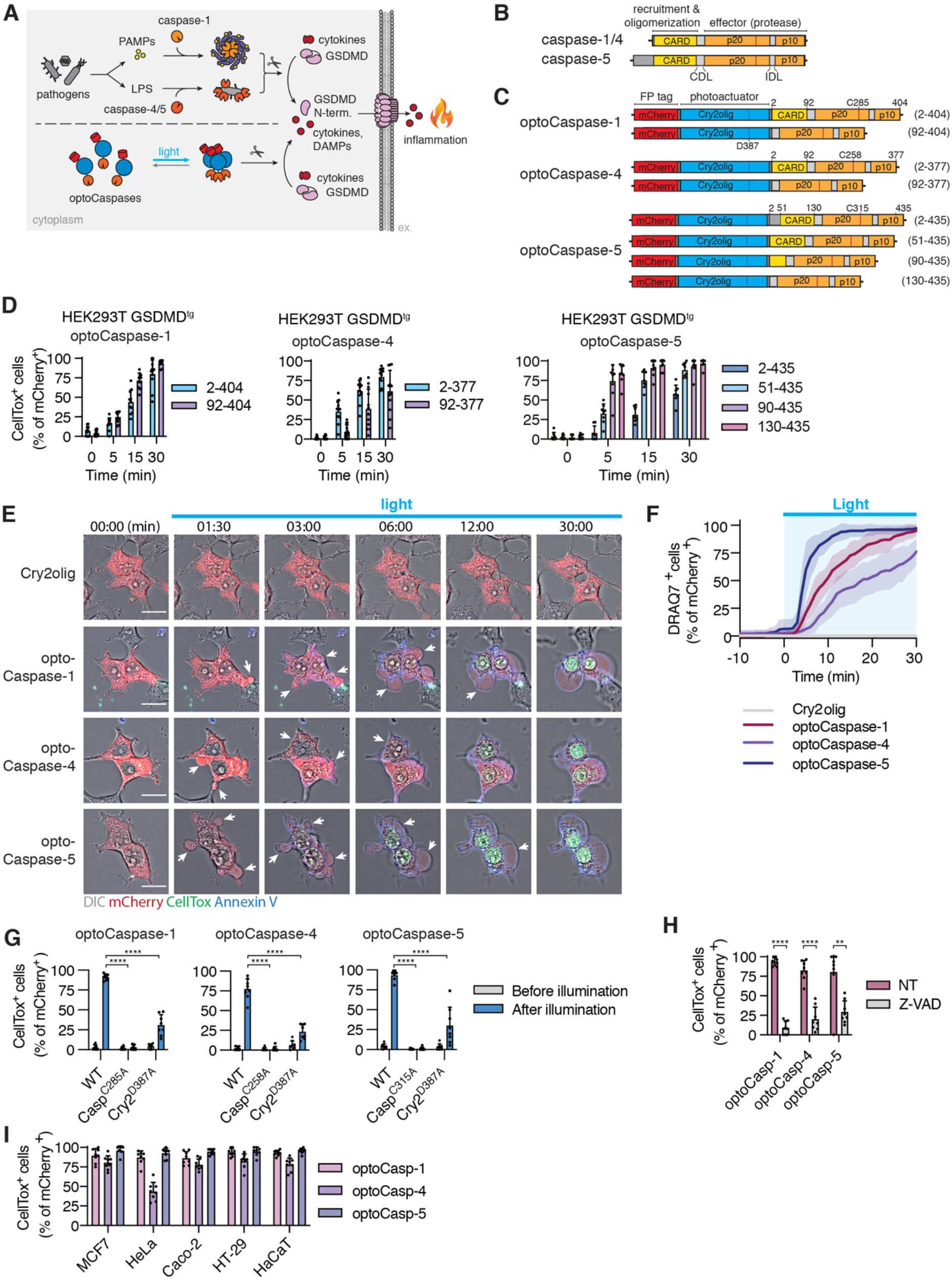
Development and validation of optogenetically activated human inflammatory caspases. **A,** Schematic representation of the inflammasome pathway and Cry2-mediated clustering of inflammatory caspases. **B,** Domain architecture of native human inflammatory caspases. **C,** Design of optoCaspase constructs. **D,** Quantification of CellTox-positive (permeabilized) GSDMD-transgenic (GSDMD^tg^) HEK293T cells expressing different optoCaspases at indicated timepoints following illumination with blue light (488nm, 5 mW/cm^2^) every 30 sec. Here and after, only cells expressing the constructs at detectable levels (mCherry^+^) at t=0 were quantified. **E,** Representative time-lapse images of GSDMD^tg^ HEK293T cells subjected to light-induced activation of Cry2olig or optoCaspase-1, −4 or −5 in presence of Annexin V (to visualize PS exposure) and CellTox Green (membrane permeabilization marker). Scale bar, 20 *µ*m. **F,** GSDMD^tg^ HEK293T cells expressing indicated constructs were imaged for 10 min in absence of blue light stimulation, and then illuminated as described in 1D for additional 30 min. The percentage of pyroptotic cells was determined at 1 min intervals by quantifying DRAQ7 (far-red membrane-impermeable DNA-binding dye)-positive nuclei. **G-H,** Percentage of CellTox^+^ GSDMD^tg^ HEK23T cells expressing wild-type or mutant optoCaspases or treated with the pan-caspase inhibitor Z-VAD-fmk before and after illumination. **I,** Percentage of CellTox^+^ cells in different cell lines expressing indicated constructs at 1 h post illumination. All data are shown as mean ± s.d. (D, G-I) or mean ± s.e.m. (F) and are representative of (E) or pooled from (D, F, G, H, I) three independent experiments. **** P < 0.0001, ** p < 0.01 (two-tailed t-test).

### Inflammatory optoCaspases induce efficient pyroptosis and downstream substrate processing in macrophage-like cell lines

Macrophages remain the most commonly used *in vitro* model to study inflammasome signaling and pyroptotic cell death. To validate optoCaspases in human macrophages we used U937 cells, a human pro-monocytic myeloid leukemia cell line that can be differentiated into a macrophage-like phenotype by PMA treatment^18^. We generated stable lines expressing mCherry-Cry2olig or optoCaspases under the control of doxycycline-inducible promoter to avoid spontaneous cell death during differentiation. Blue light stimulation of optoCaspase-expressing U937 macrophages induced rapid membrane permeabilization, Annexin-V acquisition and a typical pyroptotic morphology in the majority of mCherry-positive cells, while control macrophages expressing Cry2olig alone remain unaffected (**Fig. 2A-B**). To compare the efficacy of optogenetically-induced pyroptosis to pyroptosis caused by classical inflammasome triggers we assessed cell lysis, cytokine secretion and substrate processing. For these assays, cells were stimulated by “classical” canonical and non-canonical inflammasome activation versus blue light illumination in a custom-made light plate apparatus (LPA)^19^, which unlike confocal microscopy allows simultaneous illumination of large numbers of cells in culture plates. Nigericin (NLRP3 inflammasome activator) treatment induced around 20% of LDH release and IL-1β secretion in WT (non-transduced), mCherry-Cry2olig- and optoCaspase-expressing U937 macrophages (**Fig. 2C**). By contrast, blue light illumination efficiently killed approximately 50% of optoCaspase-1-expressing cells within 1 hour but did not induce LDH or IL-1β release from WT or Cry2olig-expressing cells. The degree of cell lysis correlated with the percentage of mCherry-positive cells in the population (**Supplementary Fig. 3A-B**), confirming that, similarly to our previous microscopy-based observations, a short illumination time was sufficient to induce the death of most optoCaspase-1-expressing cells. Immunoblot analysis confirmed optoCaspase-1 processing after blue light illumination, but not after Nigericin treatment (**Fig. 2D**), validating that the fusion protein was functioning orthogonally to the endogenous inflammasome pathway. Consistent with the LDH and IL-1β data, we observed that optoCaspase-1 efficiently processed pro-IL-1β and GSDMD **(Fig. 2D**). In line with the rapid kinetics of optoCaspase-1 activation, we found that ∼50% of optoCaspase-4 or −5-expressing cells died within 1-3 h of blue light illumination (**Fig. 2E**), whereas activation of the non-canonical inflammasome by LPS transfection required 8 h to induce even moderate cell death levels. Light-induced activation of optoCaspase-4/5 also resulted in caspase auto-processing and efficient GSDMD cleavage (**Fig. 2F**).

**Figure 2.**
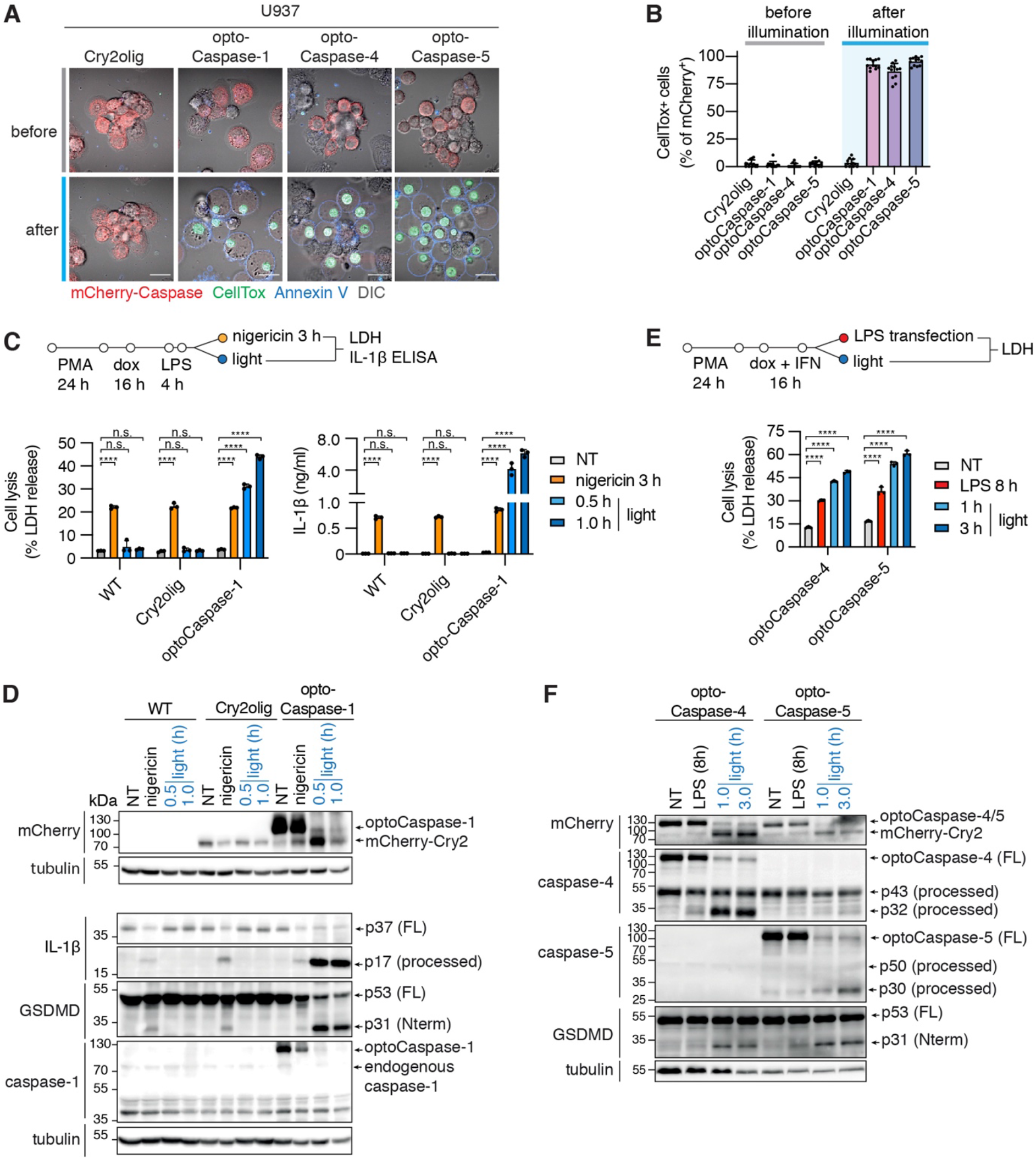
Optogenetic pyroptosis induction in human macrophage-like cell lines. **A-B,** Representative images and quantification of CellTox^+^ PMA-differentiated U937 cells expressing the indicated constructs before and after 1h of blue light stimulation (5 mW/cm^2^ every 30 sec). **C, E,** Release of LDH and IL-1β from U937 cells expressing the indicated constructs and treated with nigericin (C), transfected with LPS (E) or illuminated with blue light (4 mW/cm^2^) for the indicated time points. Non-transduced wild-type U937 were used as a control. NT, non-treated. **D, F,** immunoblot analysis of the combined cell lysates and supernatants obtained from C or E, respectively. Anti-mCherry antibodies were used to visualize full-length and processed optoCaspase-1/-4/-5 and Cry2olig. A, C-F are representative of at least 3 independent experiments, B is pooled from 3 experiments. Mean ± s.d., **** P < 0.0001, n.s. – non-significant (one-way ANOVA).

As mouse models are commonly used to study inflammasomes both *in vitro* and *in vivo*, we also generated optogenetically-activatable mouse inflammatory caspases, e.g. optoCaspase-1 and optoCaspase-11 (**Supplementary Fig. 3C**). Like human optoCaspases, mouse optoCaspase-1 and optoCaspase-11 induced high levels of pyroptosis in mGSDMD-transgenic HEK293T cells (**Supplementary Fig. 3D-E**), and their ability to induce pyroptosis was completely abrogated by mutating the catalytic cysteines. OptoCaspase-1 and optoCaspase-11 activation induced pyroptotic morphology and membrane permeabilization (as assessed by Annexin V and CellTox Green co-staining) in mouse macrophage-like RAW264.5 cells and primary murine bone marrow derived macrophages (BMDMs) (**Supplementary Fig. 3F-G**). Thus, we show that optoCaspases can be efficiently utilized both in human and mouse myeloid cell models and yield a downstream response comparable to or stronger than regular inflammasome activators.

### Sub-lytic activation of optoCaspases by limited illumination

A major advantage of optogenetics is the ability to precisely control the level of protein activation, deactivation or localization by manipulating spatial and temporal illumination parameters (such as light intensity, total illumination duration, number of light pulses, or the size of the illuminated region), and its superior reversibility upon ceasing illumination. To assess how these parameters affect optogenetic pyroptosis induction, we imaged optoCaspase-1^tg^ cells under a variety of blue light illumination conditions. Gradual increase in 488 nm light intensity resulted in a concomitant increase in the speed by which cells underwent pyroptosis (percentage of DRAQ7-positive cells over time), as well as the total number of pyroptotic cells 25 min after the beginning of illumination (**Fig. 3A**). A similar dose-response was observed in cells transiently illuminated with increasing numbers of light pulses at a single timepoint (**Fig. 3B**), and in optoCaspase-1^tg^ U937 macrophages in which increasing blue light intensity or illumination time increased the levels of cell death induction (**Fig. 3C-D**).

**Figure 3.**
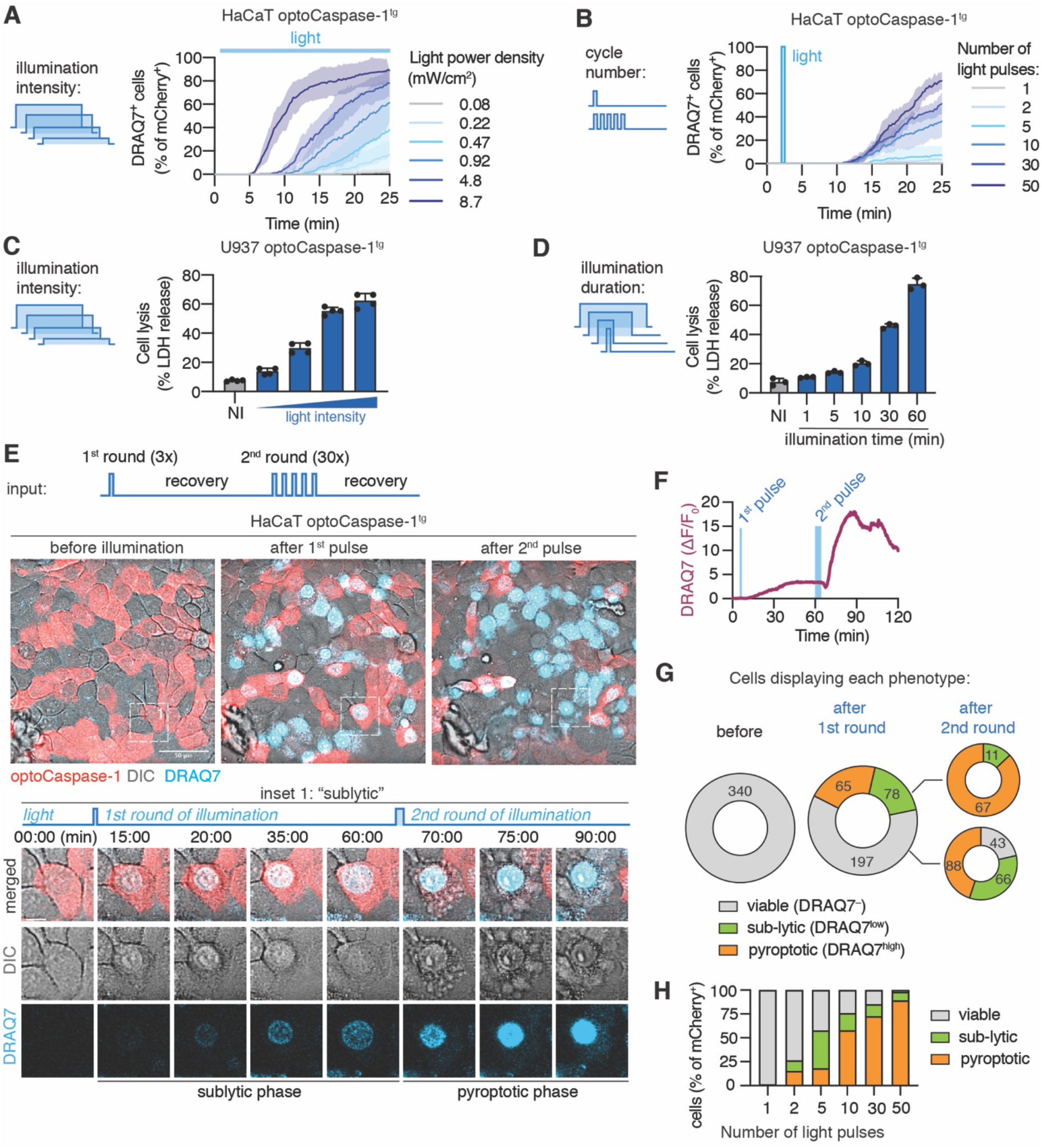
Illumination parameters determine the efficiency of optoCaspase-1-induced pyroptosis. **A-B,** OptoCaspase-1^tg^ HaCaT cells were subjected to repeated illumination with blue light of varying intensity every 15 seconds (A), or transiently illuminated at t=3 min with different number of light pulses of same intensity of 0.2 mW/cm^2^ (B) using confocal microsocpy, and percentage of pyroptotic cells was determined by quantifying DRAQ7-positive nuclei. N = 3 independent experiments, performed in triplicates, Mean ± s.e.m. from triplicate repeats **C-D,** LDH release from PMA-differentiated optoCaspase-1^tg^ U937 cells that were either continuously illuminated with light of different intensity (0.1-4 mW/cm^2^) (C) or illuminated with a defined light intensity (0.9 mW/cm^2^) for different time (D) using a Light Plate Apparatus. Representative of 3 independent experiments, Mean ± s.d. **E-H,** OptoCaspase-1^tg^ HaCaT cells were subjected to a single round of low-intensity (3 pulses x 0.2 mW/cm^2^) blue light illumination and left to recover for 60 min, after which an additional round of more prolonged illumination (10 x 0.2 mW/cm^2^) was applied. After each light stimulation, cells were classified in three distinct categories based on DRAQ7 staining: “viable” – no DRAQ7 influx and normal morphology; “sub-lytic” – low DRAQ7 influx, initial appearance of pyroptotic features (nuclear rounding, membrane blebbing) but reversion to normal morphology within 15-30 min; “pyroptotic” – high DRAQ7, pyroptotic morphology. **E,** Representative confocal images of the population of the cells displaying each phenotype before, after 1^st^ (t = 60 min) and after 2^nd^ (t = 120 min) illumination dose. Inset indicates the cell undergoing the sub-lytic pyroptosis induction and reversion after the first pulse and complete pyroptosis after the 2^nd^ pulse. Scale bar: 50 *µ*m. **F,** Normalized nuclear DRAQ7 intensity of the cell from inset 1 during sub-lytic and lytic phases of pyroptosis induction). **G,** Quantification of the cells displaying each phenotype before and after 1^st^ and 2^nd^ dose of blue light. **H,** Percentage of viable, sub-lytic and pyroptotic cells depending on the initial number of light pulses (t = 30 min after light stimulation). Data are pooled from 3 independent experiments (A, G and H) or are representative from 3 independent experiments (C-D).

Interestingly, we noticed that upon illumination with a single pulse of moderate intensity light, a fraction of cells displayed signs of early pyroptosis (e.g. membrane blebbing) and moderate DRAQ7 influx (DRAQ7^low^), but were able to survive and revert to normal morphology within 30-40 min post illumination (**Fig. 3E**, inset, and **Supplementary movie 2**). By contrast, other cells showed either no membrane permeabilization (viable nonresponding cells, DRAQ7^-^) or complete permeabilization (fully pyroptotic cells, DRAQ7^high^), the latter being followed by acquisition of typical pyroptotic morphology **(Supplementary Fig. 4A)**. Since the recovering cells (referred to as sub-lytic or DRAQ7^low^ cells) remained viable over a prolonged period of time (at least 6 hours after illumination), they were able to undergo normal cell division (with both daughter cells retaining slight DRAQ7 staining) (**Supplementary Fig. 4B**). The limited levels of DRAQ7 uptake suggest that the plasma membrane of these cells was temporarily permeabilized by GSDMD pores, but that the cells afterwards repaired the damage as had been observed for necroptosis and pyroptosis previously^20,21^. When a second round of 10 times more intense stimulation was applied, the majority of these recovered cells succumbed to complete membrane permeabilization and pyroptotic cell death (**Fig. 3F**), confirming that they retain the ability to transition into the lytic stage, and that their survival of the first pulse of illumination is not due to intrinsic defects in pyroptosis execution.

To quantitatively assess the ability of these cells to survive the transient optoCaspase-1 activation, we determined the percentage of DRAQ7-negative (live), DRAQ7^low^ (‘sub-lytic’) and DRAQ7^high^ (pyroptotic) cells. Around 40% of optoCaspase-1 cells responded to the first pulse by gaining nuclear DRAQ7 staining, and that among those around 50% underwent pyroptosis while 50% survived this low-level illumination (**Fig. 3G**). A second round of more prolonged light stimulation then induced permeabilization in 80% of cells, and among these 75% cells underwent pyroptosis and 25% survived. The second laser pulse affected DRAQ7^low^ cells more strongly than the DRAQ7^-^ cells, since a higher percentage of DRAQ7^low^ cells underwent pyroptosis after illumination. Consistent with the amount of caspase activation determining whether a cell survived or died, we observed that the fraction of “sub-lytic/DRAQ7^low^” cells was more prominent under low stimulation conditions (1-5 light pulses), yet some were present even after high-intensity stimulation (30-50 pulses) (**Fig. 3H**).

In summary, we found that caspase activation can lead to either sub-lytic or lytic GSDMD pore formation, as hypothesized before^11^, and that the optogenetic control of caspase activity yields a unique approach to study the features and mechanisms that allow cells to avoid or revert from cell death induction, such as expression of pro-survival factors or plasma membrane repair^20,21^.

### Optogenetic pyroptosis induction enables precise single-cell ablation in 2D and 3D cell culture

Laser-induced single cell ablation has been previously used to study the response of direct neighbors to dying epithelial cells^22^. To test whether optoCaspase activation enabled precise single-cell pyroptosis induction and cell ablation in populations of closely attached epithelial cells, we transiently illuminated 10-15 *µ*m^2^ regions of individual optoCaspase-1^tg^ HaCaT cells within confluent monolayers in 15-minute intervals (**Fig. 4A**). Illuminated cells underwent pyroptosis within minutes (based on morphological changes and the acquisition of nuclear DRAQ7 staining) (**Fig. 4A-B and Supplementary movie 3**), while neighboring cells remained viable and maintained membrane integrity (**Supplementary Fig. 5A**) despite similar optoCaspase-1 expression levels. The neighboring cells responded to the pyroptotic events by rapidly migrating towards the pyroptotic cell, extruding it from the monolayer and resealing of the gap (**Fig. 4A, C**).

**Figure 4.**
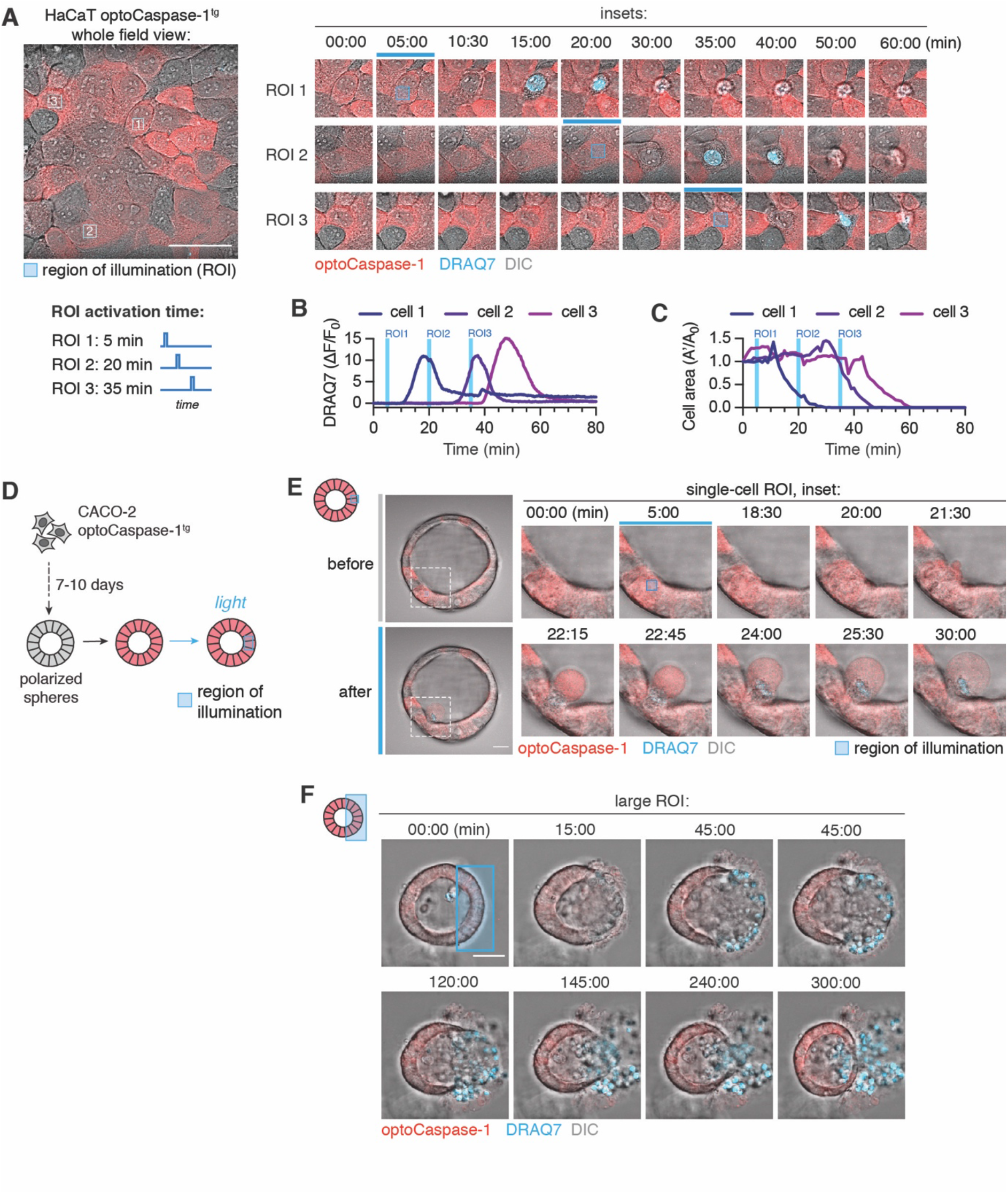
Precise optogenetic single-cell ablation in 2D and 3D cell culture conditions. **A,** Representative time-lapse images of single-cell ablation in confluent monolayer of optoCaspase-1^tg^ HaCaT cells. Left, whole field of view. Right, inset images of three selectively ablated cells; cell 1 was transiently illuminated with blue light at 5 min; cell 2 was illuminated at 20 min, and cell 3 at 35 min. Blue squares indicate the regions of interest (ROI) which were stimulated with blue light at the indicated time points. DRAQ7 (turquoise) was used to visualize membrane permeabilization during pyroptosis. **B-C,** normalized DRAQ7 intensity (B) and area of the stimulated cells from A. Vertical lines (blue) indicate the time for ROI stimulation for each cell. **D,** Schematic representation of experimental setup for spatially resolved cell ablation in 3D cultures. OptoCaspase-1^tg^ Caco-2 cells were cultured in Matrigel to induce formation of polarized spheres, and optoCaspase-1 expression was induced by doxycycline treatment. Cell death was induced by transient illumination of selected ROIs, as in A. **E-F,** representative time-lapse images showing selective ablation of a single cell (E) or large group of cells (F) in the Caco-2 spheres. Selected regions (blue squares) were transiently stimulated with blue light (3.4 mW/cm^2^) at t=5 min, and pyroptosis was determined by DRAQ7 influx. All data are representative of at least 3 independent experiments.

We next tested the optogenetic pyroptosis induction in a more complex 3D setting, using optoCaspase-1^tg^ Caco-2 cells that were cultured in a 3D microenvironment to induce the formation of polarized acini-like structures (spheres). We selectively illuminated ROIs of variable size within the sphere wall (**Fig. 4D**) and monitored cell death and neighboring cell response by time-lapse microscopy. As in 2D cell cultures, illumination of single cells within the sphere wall induced their pyroptosis, highlighted by membrane blebbing and DRAQ7 influx, and the extrusion of illuminated cells into the sphere lumen i.e. the apical side (**Fig. 4E and Supplementary movie 4**), followed by immediate gap closure by the neighboring cells. Unexpectedly, a defined luminal space was not required for extrusion to the apical side, since pyroptotic cells were still extruded towards the spheroid center in non-lumenized spheroids, which resulted in the rapid formation of a lumen around them (**Supplementary Fig. 5B).** When larger subpopulations of the cells were killed simultaneously (**Fig. 4F**), pyroptotic cell extrusion and resealing still occurred, but required a directed migration of the viable neighboring cells towards the lesion, which physically pushed out the dead cells into the lumen. Lumen size appeared to be a limiting factor for apical extrusion. Once the lumen was completely filled with dead cells, extrusion and gap closure could not proceed properly (**Fig. 4F**), which was ultimately followed by a breach in the basal membrane permitting some of the pyroptotic cells to be released from the sphere and the neighbors to reseal the spheroid.

In summary, these data demonstrate that optogenetic caspase activation can be used for a precise single cell ablation in 2D and 3D cell culture models and highlight the immense potential of the new toolset for the study of bystander cell responses and cell extrusion in these models.

### Optogenetic induction of apoptosis

The lytic cell death caused by pyroptosis is in stark contrast to apoptosis, an immunologically silent form of PCD. Extrinsic apoptosis is induced by signaling complexes such as the DISC or TNFR1 complex IIa/b, which recruit and activate the initiator caspase-8 via homo-typic DED-DED interactions (**Fig. 5A**). In the intrinsic apoptotic pathway, release of cytochrome c from mitochondria triggers the assembly of the apoptosome that recruits and oligomerizes caspase-9 through CARD-CARD interactions. Both initiator caspases cleave and activate the executioner caspases-3/-7 to drive cell death. To expand our toolset to optogenetic apoptosis induction, we fused Cry2olig N-terminally to either full-length or ΔCARD/ΔDED caspase-8/-9 (**Fig. 5B**). Illumination of optoCaspase-8- or optoCaspase-9-expressing HEK293T cells induced appearance of classical apoptotic features, such as cell body shrinking, surface detachment and extensive membrane blebbing in the majority of mCherry-positive cells within 30 min (**Fig. 5B-C, Supplementary Fig 6A and Supplementary movie 5**). Since the DEDs of caspase-8 and to a lesser extent the CARD of optoCaspase-9 delayed apoptosis induction (**Fig. 5C**), we used ΔCARD/ΔDED optoCaspase-8/-9 for further characterization of the constructs.

**Figure 5.**
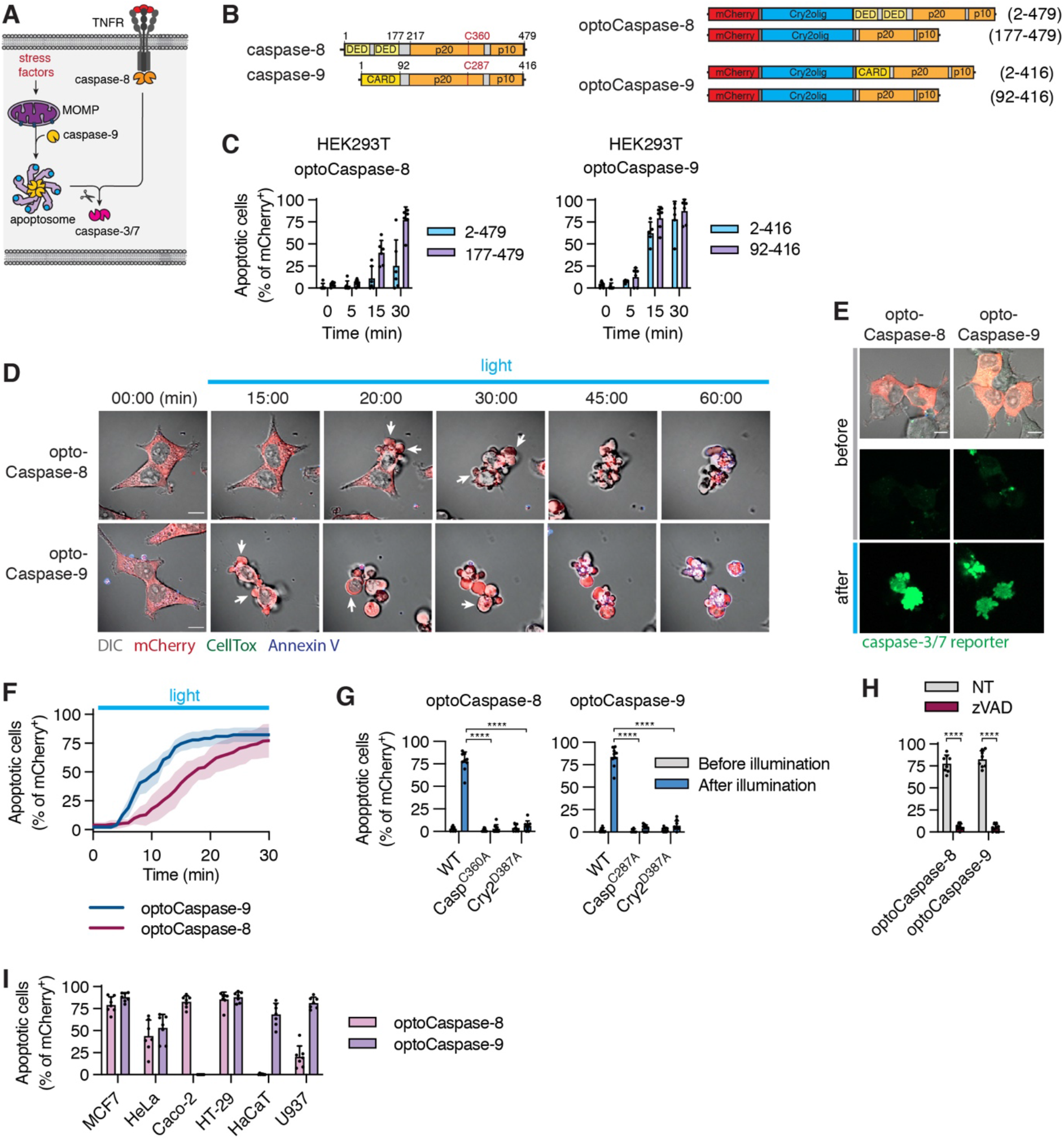
Optogenetic activation of apoptotic initiator caspases. **A, B,** schematic representations of intrinsic and extrinsic apoptosis induction (A) and of full-length or truncated optoCaspase-8 and optoCaspase-9 (B). **C,** HEK293T cells expressing different optoCaspase-8/9 versions were subjected to blue light illumination (5 mW/cm^2^), and percentage of apoptotic cells over time was determined based on cell morphology (cell shrinking, blebbing, nuclear fragmentation). **D,** representative time-lapse images of HEK293T cells expressing DED-deficient optoCaspase-8 and CARD-deficient optoCaspase-9 undergoing light-induced apoptosis (5 mW/cm^2^). Annexin V (blue) was used to visualize PS exposure. Scale bar, 10 *µ*m. **E,** Representative images of caspase-3/7 reporter (green) activation in optoCaspase-8/9-expressing HEK293T cells before and after 1 h of blue light stimulation (5 mW/cm^2^). **F,** dynamics of optoCaspase-8 and optoCaspase-9-induced apoptosis. Cells were stimulated with blue light as in C, and the percentage of apoptotic cells was quantified every min. **G-I**, Percentage of cells expressing wild-type (G, I) or mutated (G) optoCaspases or treated with Z-VAD (H) and displaying apoptotic morphology before and 1 h after light stimulation (5 mW/cm^2^). All data are representative of (D) or pooled from (C, E-I) three independent experiments. P < 0.0001 (two-tailed t-test).

Validation of optoCaspase-8/-9 showed rapid activation of a genetically encoded caspase-3/-7 activity reporter confirming efficient activation of executioner caspases, which preceded acquisition of Annexin-V staining (**Fig. 5D-F, Supplementary Fig. 6B**). Catalytic dead or dimerization-deficient mutant constructs failed to induced cell death (**Fig. 5G**). Treatment with the pan-caspase inhibitor Z-VAD abrogated optoCaspase-8/9-induced apoptosis (**Fig. 5H**). Finally, we tested the ability of optoCaspase-8/-9 to induce apoptosis in different cell lines (**Fig. 5I**). We observed a strong variation in optoCaspase-8 and optoCaspase-9 induced apoptosis levels between cell lines (**Fig. 5I**), potentially due to variable levels of endogenous caspase inhibitors (cIAPs/XIAP). Consistent with this notion, treatment with the SMAC mimetic AZD5582 facilitated apoptosis induction in at least some cell lines (**Supplementary Fig. 6C**). The inhibitor did not affect optoCaspase-8-induced apoptosis in HaCaTs or optoCaspase-9-induced apoptosis in Caco-2, suggesting that other mechanisms, such as inhibitory caspase phosphorylation, might block initiator caspase activation. In summary, optogenetically-activatable caspase-8/-9 can be used to induce apoptosis in a variety of cell lines and might provide a new approach to study endogenous regulatory mechanism that control apoptosis execution.

### Optogenetic activation of necroptosis by Rip3 and MLKL

To complete the toolbox of optogenetic cell death inducers, we next focused on necroptosis, a form of cell death that is often considered to be a back-up pathway to apoptosis. Under conditions where the activity of caspase-8 is blocked, activation of TNFR1 or TLRs triggers the formation of the necrosome, a signaling platform that engages a kinase signaling cascade via RIP1/3 culminating in phosphorylation and oligomerization of the death executioner MLKL (**Fig. 6A**). To optogenetically induce necroptosis, we constructed C- or N-terminal fusions of mCherry-tagged Cry2olig to wild-type or RHIM (RIP homotypic interaction motif)-deficient (QIG449–451AAA mutation) full-length human RIP3 protein (**Fig. 6B**), or the NBB-PKD (N-terminal bundle and brace-protein kinase domain) of MLKL. Illumination of optoRIP3 or optoMLKL-expressing HEK293T cells resulted in cell rounding and PS exposure (Annexin-V staining), indicating that cell death had been induced (**Fig. 6 C-D**). Among the four RIP3 constructs, the N-terminal Cry2 fusion to optoRIP3ΔRHIM (in the following termed optoRIP3) proved the most efficient in inducing cell death (**Fig. 6C**), while the other constructs proved less efficient or led to dark-state protein aggregation and spontaneous apoptosis-like cell death in a significant fraction of mCherry-positive cells (**Supplementary Fig. 7A-C**).

**Figure 6.**
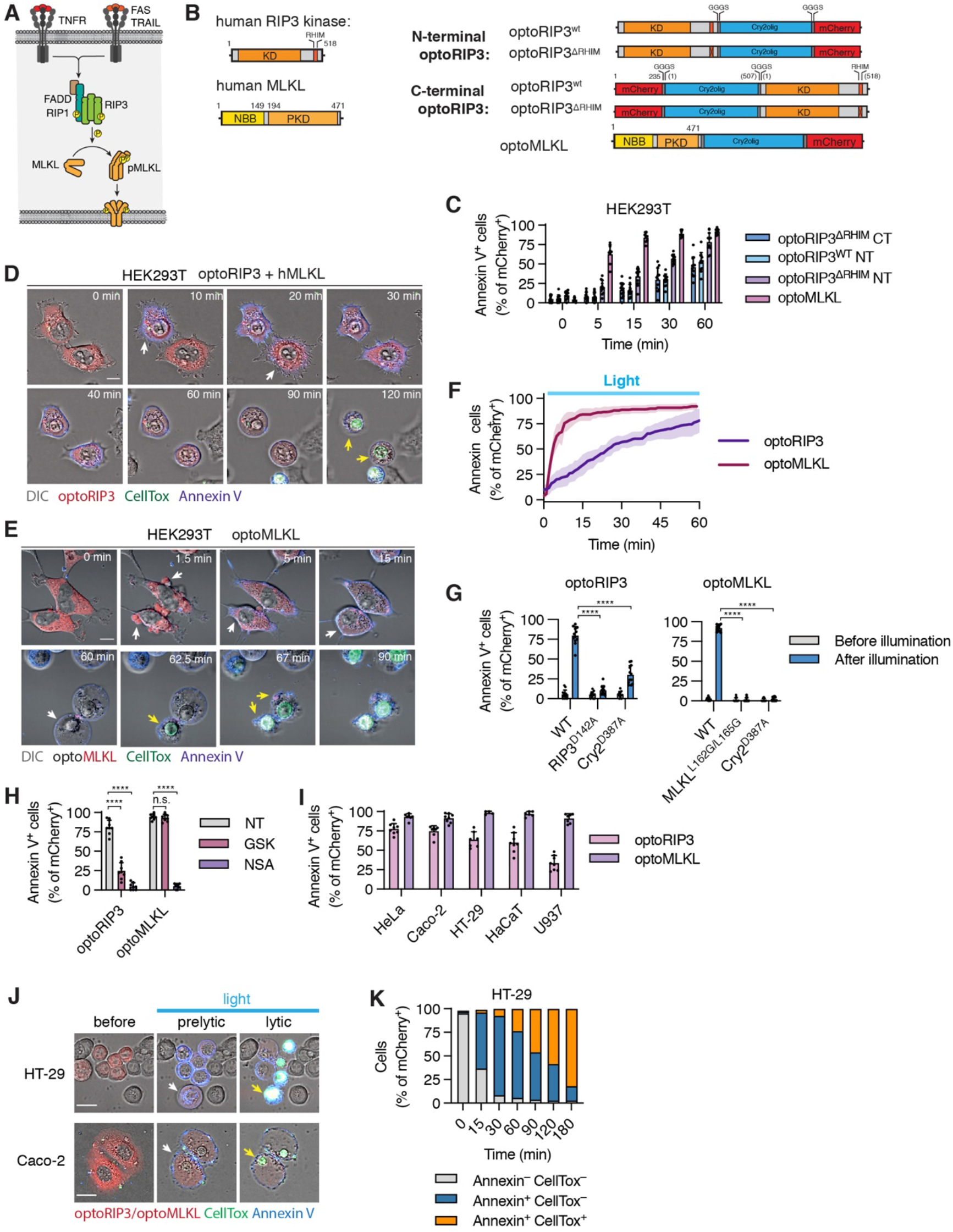
Optogenetic activation of necroptotic effectors optoRIP3 and optoMLKL. **A,** Schematic illustration of the necroptosis pathway. **B,** Domain architecture of native human RIP3 and MLKL proteins and designed of optoRIP3 and optoMLKL constructs. **C,** Quantification of Annexin-positive HEK293T cells expressing different optoRIP3 versions or optoMLKL at indicated timepoints. Cells were photostimulated with 488 nm laser (5 mW/cm^2^) every 15-30 sec. Due to the lack of endogenous MLKL expression in HEK293T, optoRIP3 constructs were co-transfected with human MLKL. Due to the notable delay between PS exposure (early necroptosis) and membrane permeabilization (late necroptosis), hereafter Annexin-V staining was utilized as a primary readout for optoRIP3/optoMLKL activation. **D-E,** representative time-lapse images of HEK293T cells undergoing light-induced necroptosis upon optoRIP3 (N-terminal, RHIM-deficient) or optoMLKL activation. Annexin V (blue) is used to visualize PS exposure during early (non-lytic) necroptosis phase, and CellTox staining (green) marks the loss of membrane integrity during late (lytic) stage. **F,** Dynamics of necroptosis induction via optoRIP3 and optoMLKL activation. **G-H,** Cells expressing wild-type, catalytically deficient or oligomerization-deficient optoRIP3/optoMLKL (G) or treated with 1 *µ*M GSK’872 (RIP3 inhibitor) or 5 *µ*M necrosulfonamide (MLKL inhibitor) were subjected to blue light illumination for 1 h. **I,** Necroptosis induction in various human cell lines. Percentage of AnnexinV+ cells was assessed after 1 h of continuous illumination (5 mW/cm^2^ every 15 sec). **J,** representative confocal images of non-stimulated, early (pre-lytic) and late (permeabilized) necroptotic Caco-2 and HT-29 cells expressing optoMLKL. **K,** Quantification of optoMLKL-expressing HT-29 cells undergoing light-induced necroptosis and displaying only Annexin V+ (early necroptosis) or Annexin V+ CellTox (late necroptosis) staining. Data are representative (D, E, J) or pooled from (C, F-I and K) 3 independent experiments, mean ± s.d., **** p < 0.0001, n.s. – non-significant (two-tailed t-test).

Time-lapse microscopy of HEK293T cells expressing either optoRIP3 (co-expressed with hMLKL) or optoMLKL revealed that constructs induced a rapid necrotic cell death upon blue light illumination, that was characterized by cell rounding, Annexin-V staining and at later stages by CellTox acquisition (**Fig. 6D-E and Supplementary movie 6**). Direct kinetic comparison of necroptosis induction over time revealed that while both constructs induced the death of the majority of mCherry-positive cells within 1 hour of illumination, optoRIP3-induced necroptosis was slower compared to optoMLKL **(Fig. 6F)**, suggesting that additional factors, such as endogenous regulation of RIP3 or kinetics of MLKL phosphorylation and assembly, may contribute to this difference. OptoRIP3 did not induce any phenotypical changes in absence of hMLKL co-expression **(Supplementary Fig. 7D)**, confirming that morphological changes and cell death resulted from RIP3-dependent MLKL phosphorylation. Mutation of RIP3 D142 or MLKL L162/L165 significantly decreased the proportion of necroptotic cells upon illumination (**Fig. 6G**), as did the treatment with RIP3 or MLKL inhibitors GSK’872 and necrosulfonamide (NSA) **(Fig. 6H).** Both constructs were highly active in a range of cell types commonly used for necroptosis studies (**Fig. 6I**).

Interestingly, morphological analysis revealed that necroptosis proceeded through two distinct phases. An initial ‘sub-lytic’ phase of approximately one hour, during which cells displayed membrane blebs, acquired Annexin-V staining and rounded up, but maintained membrane integrity, and a lytic phase characterized by sudden and complete rupture and disappearance of the cell envelope and rapid CellTox acquisition **(Fig. 6D-E and J-K and Supplementary movie 6)**. This was strikingly different from pyroptotic cells which usually acquired CellTox staining already during the blebbing and ballooning phase, and never displayed a complete rupture and disappearance of the plasma membrane (at least over the time we imaged the cell). This suggests that, during pyroptosis, GSDMD pore formation and membrane permeabilization is an early event, while membrane permeabilization was a late event during necroptosis and was preceded by a sub-lytic phase during which MLKL channels are formed, but do not yet reach the threshold induce full PMR as suggested previously^23^. In conclusion, these studies not only validate the usefulness of optogenetic tools for single cell studies, but also highlight the unique features of individual cells when progressing into different types of cell death.

### Optogenetic activation of zebrafish caspases allows spatial and temporal controlled induction of pyroptosis and apoptosis *in vivo*

Zebrafish (*Danio rerio*) embryos are a powerful model to visualize biological processes like cell differentiation and death in real-time and *in vivo*. Zebrafish feature homologs of mammalian apoptotic and inflammatory caspases, such as caspa which induces pyroptosis in zebrafish skin cells downstream of ASC-dependent inflammasomes^24^, and caspb that assembles a non-canonical inflammasome together with zfNLRP3^25^ (**Supplementary Fig. 8A**). Similarly to mammalian optoCaspases, we designed optogenetically activatable opto-zf(zebrafish)Caspa, opto-zfCaspb and opto-zfCaspase-8 by replacing their PYD or DED by Cry2olig and evaluated these constructs in GSDMD^tg^ or wild-type HEK293T cells **(Fig. 8A and Supplementary Fig. 8B)**. Activation of opto-zfCaspa/b in HEK293T cells induced rapid pyroptosis and CellTox influx, while optoCaspase-8 activation triggered the appearance of a typical apoptotic morphology (**Supplementary Fig. 8B-D**), confirming both functionality and evolutionary conservation of substrate specificity of these proteins.

We next transferred the constructs into an inducible system that allowed the expression of optogenetic constructs under control of a heat shock responsive element (HSE) ^26^. We created stable zebrafish lines by tol2-based transgenesis and used a cardiac myosin light chain 2 (cmlc2) promoter driving tagRFP expression in the heart as a way to identify transgenic larvae (**Fig. 7B**). Heat-shock treatment induced efficient opto-caspase expression, yielding a typical mosaic expression pattern in different tissues, but did not result in any detectable spontaneous cell death induction (**Fig. 7C**). While skin cells (keratinocytes and basal cells) are able to form inflammasomes and dye in response, muscle cells naturally not express ASC, inflammatory caspases or gasdermins. The mosaic expression of opto-zfCaspases in these different tissues (schematically depicted in Fig. 7D) allow us to compare the effect of different cell death stimuli in inflammatory responsive and unresponsive tissues. Blue light illumination of Opto-zfCaspa-expressing keratinocytes (**Fig. 7D-E and Supplementary movie 7**) rapidly induced pyroptosis as judged by the appearance of a characteristic pyroptotic morphology highlighted by shrinking and eventually membrane ballooning (**Fig. 7E, yellow arrows**). By contrast, opto-Caspase-8 activation in keratinocytes and muscle cells induced apoptosis, which was highlighted by the formation of apoptotic bodies (**Fig. 7F and Supplementary movie 8**). While the effect of opto-caspase induction in skin cells is visible within minutes the effect of their activation in muscle cells only becomes apparent after several hours (**7F, white arrows**). Unexpectedly, opto-zfCaspb also induced apoptosis in keratinocytes (**Supplementary Fig. 7E**) despite its ability to cleave human GSDMD in HEK293 cells. Both opto-zfCaspa as well as opto-zfCaspase-8 induced apoptotic cell death in muscle cells (**Fig. 7E and F**) visible as a contraction without balooning of the cell), possibly due to the absence of gasdermins in these cells^27^. In line with our previous observations in mammalian system, induction of pyroptosis in zebrafish skin triggered the rapid extrusion of the pyroptotic cells from the monolayer, followed by the acquisition of DRAQ7 staining **(Fig. 7G and Supplemental movie 9)**. Finally, we also used a 2-photon laser to stimulate selected regions (**Fig. 7H**, light blue squares), demonstrating that we can restrict activation of cell death to single cells. In summary, these results show that optoCDEs allows both temporal and spatial activation of cell death within living tissue down to single cell level.

**Figure 7:**
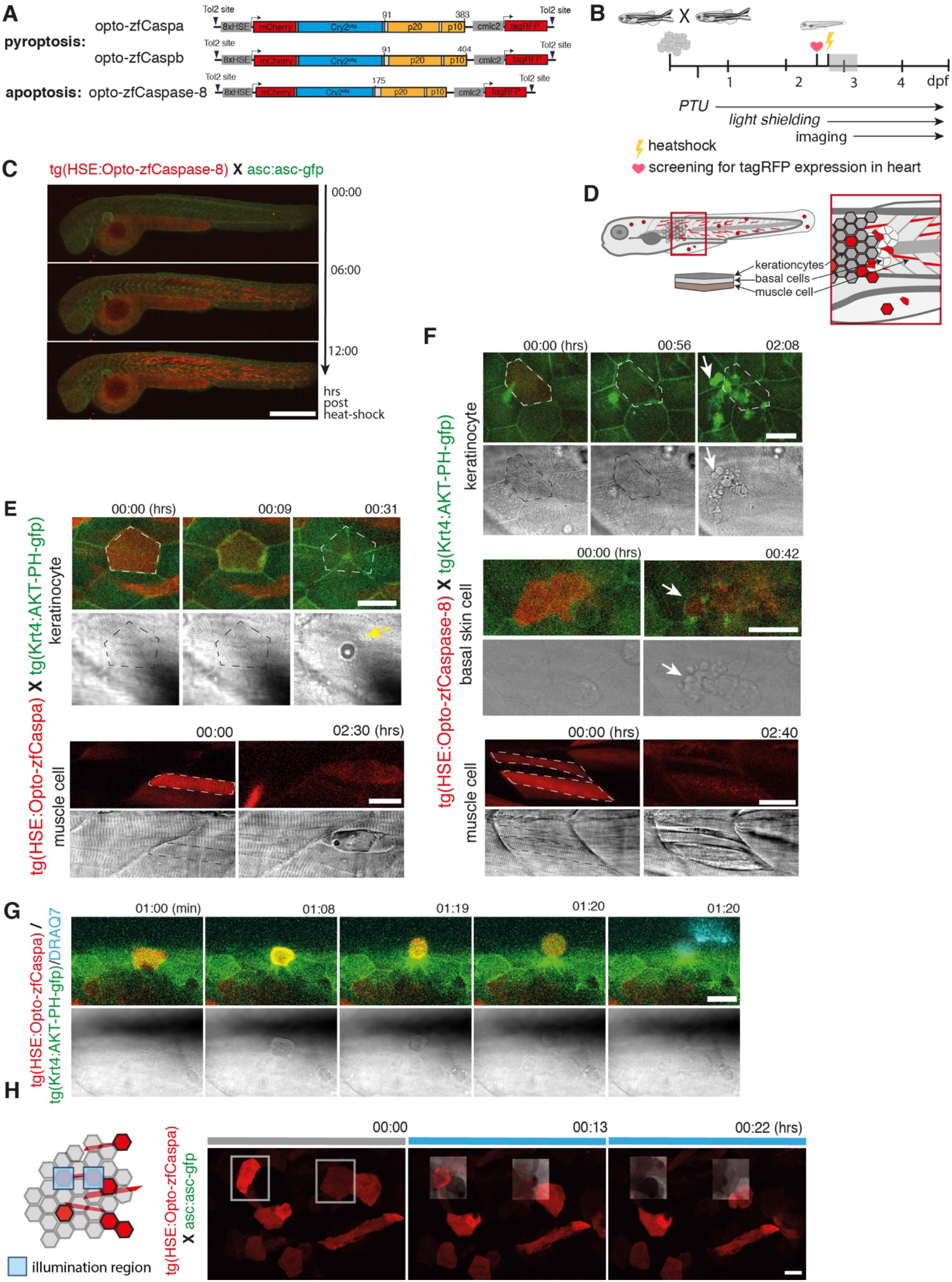
Spatial and temporal control of programmed cell death in zebrafish by Opto-caspases *in vivo*. **A**, Design of Opto-caspase construct for heat-shock-induced expression of mCherry-Cry2olig fusion with zfCaspa, zfCaspb or zfCaspase-8. Numbers depict the boundaries between different domains. Depicted are also the Tol2 site flanking the Opto-caspase cassette and the heart specific promoter cmlc2 driving tagRFP expression. **B**, Experimental setup for expression, activation and imaging of optogenetic induction of PCD. Grey shaded is the time frame in which panel C was acquired. **C**, Time-lapse imaging of 2,5-dpf Tg(Opto-caspa)Xasc:asc-gfp larvae after heat shock. Shown are time points corresponding to the time directly after heatshock, and 6 and 12 hphs. Scale bar 500 *µ*m. **D**, Schematic depiction of a zebrafish larvae showing location of keratinocytes, basal cells and muscle cells. **E-G** Opto-caspase lines were crossed to tg(Krt4:AKT-PH-GFP)(green). Opto-caspase construct expression in red. Time scale is in hours, starting at the first timepoint of 488 laser exposure. Fluorescent images are z-projections, bright field images are single planes. Dying cells are outlined by dashed lines. Scale bar for all images are 20 *µ*m. **E**. Representative time lapse images of keratinocyte pyroptosis and muscle cell death after activation of Opto-zfCaspa and single plane bright field image. **F**. Representative time lapse images of keratinocyte, basal cell and muscle cells dying after activation of Opto-zfCasp8. **G**, Representative time lapse images of keratinocyte, after activation of Opto-zfCaspb. **H**, Two photon-activation of Opto-zfCaspa (red) in single cells. Illumination frame is highlighted in grey. Opto-zfCaspa was crossed to asc:asc-gfp line, asc-gfp (green) is visible only in 2-Photon illuminated squares. The starting frame is right before 2-Photon activation. Scale bar is 20 *µ*m.

### Distinct cell death modes induce differential fates of epithelial cells

To highlight the usefulness and versatility of the novel tools for the study of complex biological questions, we compared the fates of cells dying by lytic (pyroptosis or necroptosis) versus non-lytic (apoptotic) cell death in the context of confluent epithelial monolayers. Previous studies reported that apoptosis is followed by extrusion of dead cells from monolayers, but death was usually induced non-specifically by laser ablation that causes a mixed apoptotic/necrotic phenotype^6–8^ and fates of necroptotic/pyroptotic cells have not been studied on a single cell level before.

To avoid any confounding effects of low-level cell death activation in neighboring cells due to light diffusion, we generated confluent mosaic monolayers of optoCaspase-1-, optoCaspase-8- or optoMLKL-expressing Caco-2 cells co-cultured with 20-50-fold excess of wild-type cells (**Fig. 8A**). Illumination of mCherry-positive optoCDE-expressing cells initiated the respective cell death programs, which was followed by a rapid rearrangement of neighboring cells and re-establishment of epithelial integrity (**Fig. 8B and Supplementary movie 10**). Consistent with our previous observations (**Fig. 3**), both pyroptotic and necroptotic cells were rapidly extruded from the monolayer within minutes following the appearance of the first morphological signs of cell death, Annexin-V staining and, in case of pyroptosis, CellTox influx (**Fig. 8B-E and Supplementary Fig. 9A-B**), with only a small fraction of Annexin-V-positive membrane remnants being left behind and taken up by the neighboring cells **(Supplementary Fig. 9B)**.

**Figure 8.**
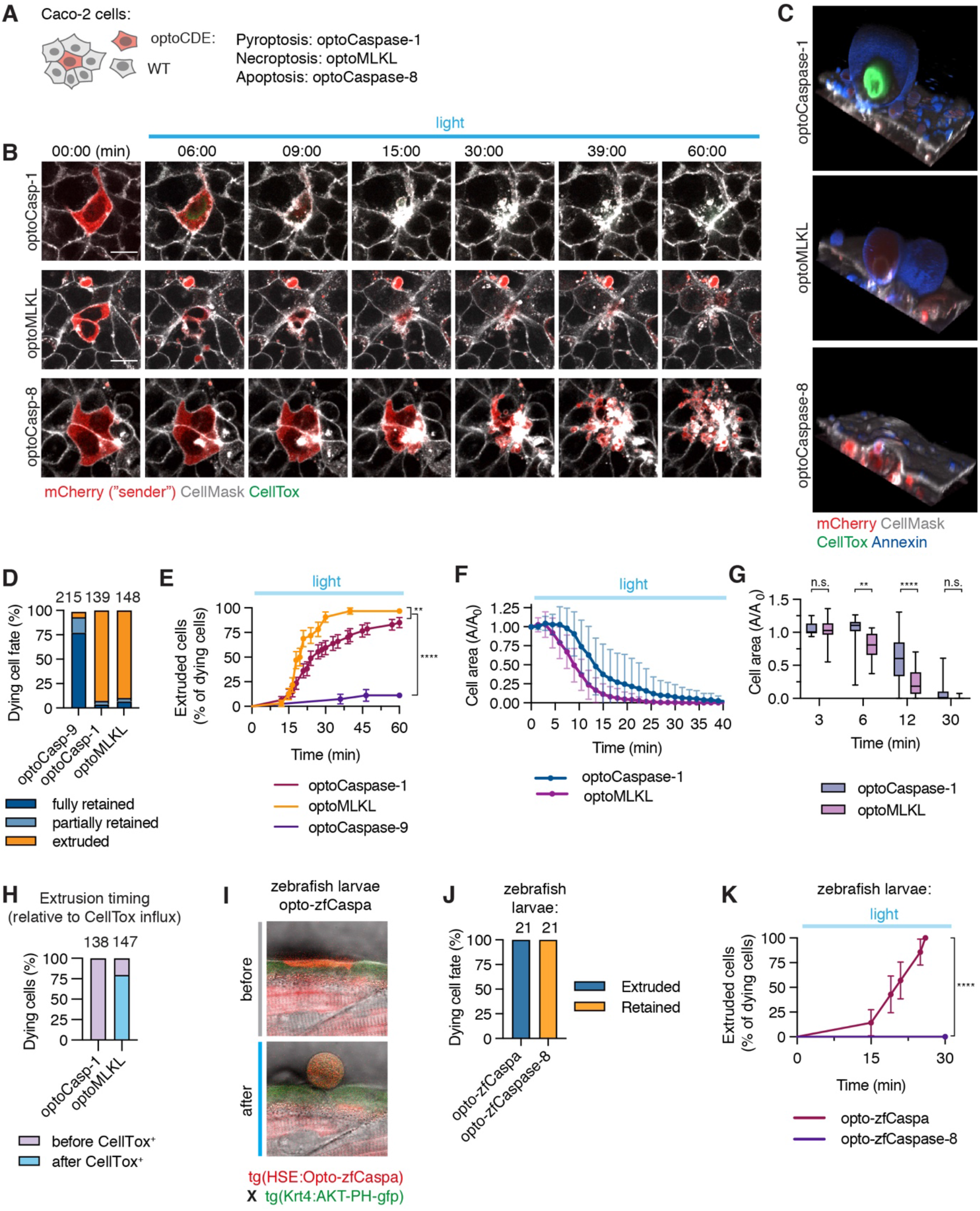
Response of neighboring cells to programmed cell death. **A,** Illustration of co-culture system consisting of “sender” cells expressing mCherry-tagged optoCDEs (red) and 50x excess number of wild-type neighbors. **B,** Time-lapse images of cells undergoing optoCaspase-1-induced pyroptosis, optoCaspase-8-induced apoptosis or optoMLKL-induced necroptosis in co-cultures. Membranes were stained with CellMask (grey) to visualize cell boundaries, and membrane permeabilization was determined by CellTox staining (green). Scale bar, 20 *µ*m. Cells were photostimulated with 488 nm laser (8.7 mW/cm^2^) every 90 sec. **C,** 3D reconstructions showing extruded pyroptotic and necroptotic cells (top and middle panels) or efferocytosed apoptotic bodies (bottom panel) at the end timepoint (1 h). Annexin V (blue) labels necrotic cell membranes. **D,** Quantification of dying cell fates. Clusters of 1 or several optoCDE-expressing cells surrounded by wild-type cells were stimulated with 488 nm light (8.7 mW/cm^2^) every 180 sec for 90 min of total time. Cells were classified as “extruded” if over 80% of cell body was expelled, “partially extruded” if between 20 and 80% of cell body was expelled, and “retained” if over 80% of cell body (including nucleus) remained within the imaging plane at the end of experiment. Numbers above bars indicate total number of cells quantified per condition (pooled from 3 independent experiments). **D,** Quantitative analysis of pyroptotic, necroptotic and apoptotic cell extrusion. To simplify the analysis, cells were classified as “extruded” if over 50% of cell body was expelled. N = 56, 32 and 27 cells pooled from 3 experiments. ** p<0.01, **** p<0.0001 (Mantel-Cox test). **E-G,** Analysis of pyroptotic and necroptotic cell area (normalized to cell area at t=0) at indicated timepoints. Mean N = 20 and 18 cells, 3 independent experiments. * p<0.05, ** p<0.01 (two-tailed t-test). **H,** Percentage of pyroptotic and necroptotic cells fully extruded before or after gain of CellTox signal, pooled from 3 independent experiments. **I,** representative images of opto-zfCaspa-expressing cell in zebrafish skin before and after stimulation with 488 nm laser**. J,** Quantification of extruded vs. retained cells during opto-zfCaspa-induced pyroptosis and opto-zfCaspase-8-induced apoptosis in zebrafish larvae. **K,** Analysis of pyroptotic and apoptotic cell extrusion in zebrafish larvae. N = 21 events for each type of cell death, **** p < 0.0001 (Mantel-Cox test).

The majority of apoptotic cells on the other hand were retained within the monolayer, where they were rapidly fragmented into apoptotic bodies and either partially or fully taken up by the neighboring cells (**Fig. 8B-E**). These apoptotic bodies persisted within neighbors for extended periods of time (up to 12 h) before being eventually degraded (**Supplementary Fig. 9C-E).** Unexpectedly, apoptotic cell fragmentation of Caco-2 cells was only seen in the presence of neighboring cells, given that apoptosis induction in isolated groups of optoCaspase-8-expressing cells did not give rise to apoptotic bodies, despite displaying other apoptotic features, such as cell shrinking and acquisition of Annexin-V signal (**Supplementary Fig. 9F-G**). Apoptotic cell fragmentation and engulfment preceded PS exposure and was not blocked by incubation with Annexin-V **(Supplementary Fig. 9F and H)**, suggesting that, unlike in case of efferocytosis by professional phagocytes, PS sensing was not a major driver of apoptotic cell engulfment by epithelial cells. Importantly, retention of cells followed by fragmentation and engulfment was also observed in cases of spontaneous apoptosis of wild-type cells **(Supplementary Fig. 9I**), confirming that engulfment was neither a consequence of laser illumination itself nor of the lack of natural apoptotic signals.

Since a mayor difference between apoptotic and necrotic cells (pyroptosis/necroptosis) is the loss of membrane integrity, we investigated if extrusion of pyroptotic and necroptotic cells correlated with membrane permeabilization. Time lapse analysis revealed that while membrane permeabilization and CellTox influx always preceded extrusion of pyroptotic cells, the majority of necroptotic cells were fully extruded even before acquiring CellTox signal, and marginally faster than pyroptotic cells (20 vs. 26 min on average) **(Fig. 8F-H and Supplementary Fig. B)**. Thus, this observation implied that extrusion is not driven by cell lysis, but by other signals stemming from pyroptotic and necroptotic cells. The same difference in the fate of dying cells (i.e. the efferocytosis of apoptotic cells vs. the extrusion of necrotic cells) was found upon optogenetic induction of different types of cell death *in vivo* upon induction of apoptosis or pyroptosis in zebrafish larvae (**Fig. 7E-G and Fig. 8I-LK**). In summary, our data demonstrate that unlike unspecific induction of mixed apoptotic/necrotic cell death modes by laser ablation, the specific cell death induction using optoCDEs allows to identify previously unknown differences in the ways the epithelia respond to apoptotic and necrotic cells.

### Extrusion and uptake of dying cells requires active cytoskeleton remodeling

Both efferocytosis or extrusion of dead cells involves extensive remodeling of the cytoskeleton and the formation of characteristic structures like phagocytic cups or contractile actomyosin rings ^22,28–30^. We thus used Lifeact-GFP-expressing Caco-2 cells to visualize actin dynamics in cells neighboring apoptotic or pyroptotic/necroptotic cells. While resting neighbors showed an actin rich cortex and dynamic randomly oriented lamellipodial protrusions at the basal side **(Supplementary Fig. 10A),** we found that neighbors involved in the extrusion of necrotic cells (pyroptotic/necroptotic) formed polarized lamellipodia at the basal cell surface (0 *µ*m, asterisk) as well as contractile purse string-like structures, visible as thick Lifeact-positive structures around the dying cell, at the apico-lateral side (arrowheads) **(Fig. 9A-B and F**). These features are consistent with the cytoskeletal structures that were previously implicated in the extrusion of laser-ablated dead cells^30,31^.

**Figure 9.**
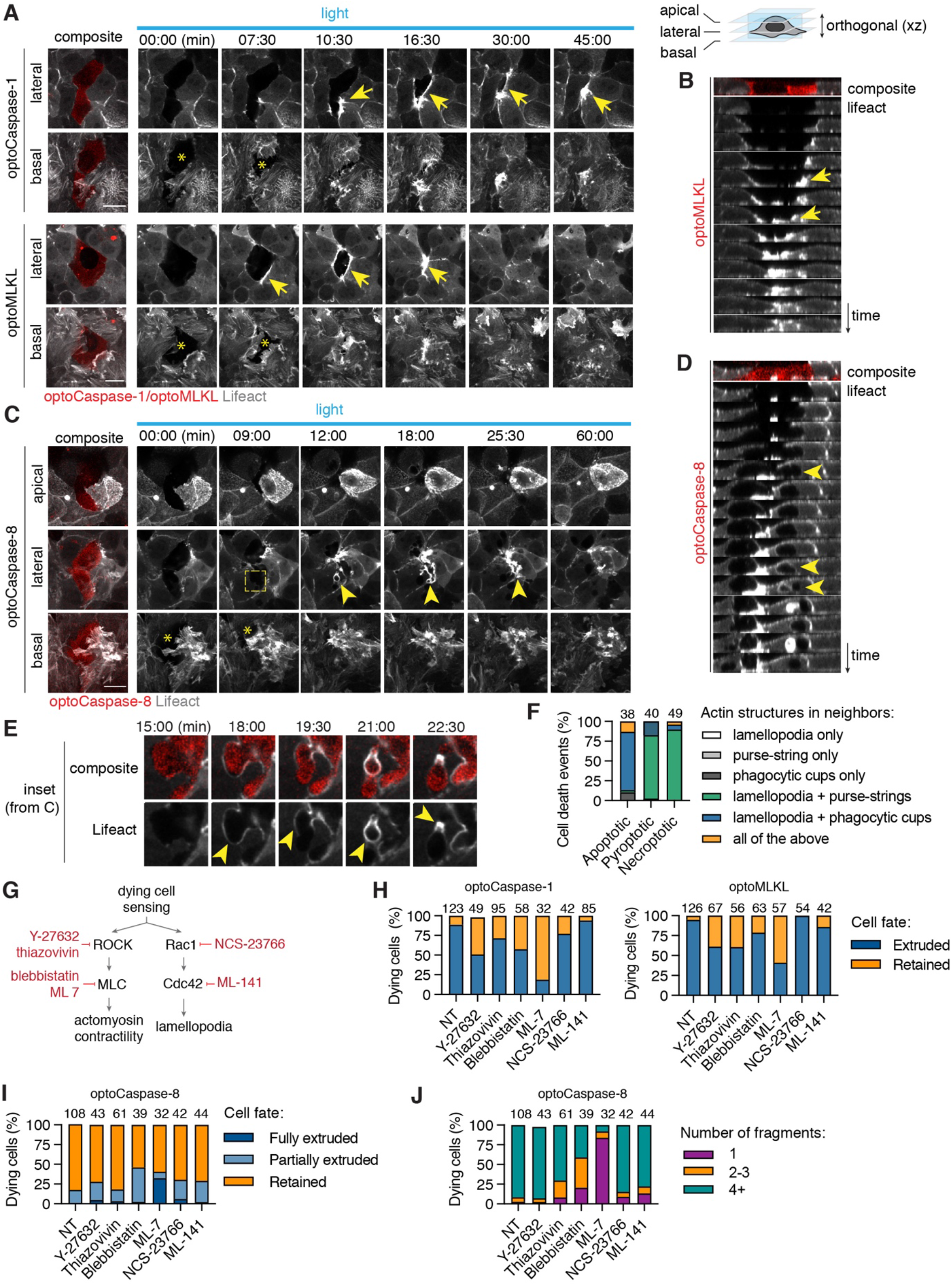
Necrotic cell extrusion and apoptotic cell efferocytosis require differential cytoskeletal rearrangement in neighbors. **A,** Time-lapse images showing actin dynamics in neighboring cells during pyroptotic and necroptotic cell extrusion from the monolayers. White arrows indicate formation of lamellipodia at the basal (0 *µ*m) plane, and yellow arrows show actin cables at the medial (+ 5 *µ*m) plane. Photostimulation (2 mW/cm^2^) and z-stack acquisition were performed every 90 sec. **C,** Time-lapse images of cytoskeletal rearrangements in neighbors during apoptotic cell fragmentation and engulfment, showing apical closure (apical plane), efferocytic cup-like structure formation (medial plane, yellow arrows) and lamellopodia (basal plane) around apoptotic cell. Cells were stimulated and imaged as in A. Apical plane corresponds to the maximum intensity projection of 4 planes acquired at 0.6 *µ*m intervals. **B, D,** reconstructed side views of cytoskeletal changes during necroptotic cell extrusion (from A) or during apoptotic cell fragmentation and engulfment (from C). **E,** Inset images from C showing actin polymerization in neighbors during apoptotic corpse fragmentation and uptake. **F,** Quantification of actin structures formed in neighboring cells in response to different types of cell death. Numbers indicate the total number of cell death events per conditions, data is pooled from 3 independent experiments. **G,** Schematic representation of the major regulators of actin remodeling during lamellipodia and purse-string formation, and inhibitors targeting them. **E-G,** Quantification of pyroptotic, necroptotic **(E)** and apoptotic **(F)** cell extrusion (at t = 60 min) and apoptotic cell fragmentation in co-cultures treated with the different cytoskeletal inhibitors. Cells were photostimulated with blue light 8.7 mW/cm^2^ every 180 sec. Data are pooled from 6 independent experiments; total number of cell death events is indicated above the bars for each condition. Scale bars: 20 *µ*m.

Apoptosis induction, on the other hand, led to a rapid polarized apical movement of the neighbors on top of the apoptotic cell, starting immediately after the appearance of the first signs of apoptosis and preceding apoptotic cell fragmentation. This was accompanied by the simultaneous engagement of basal lamellipodia (asterisk), which dynamically extended below the apoptotic cell, leading to sealing of the basal gap. Neighboring cells polymerized actin from the apico-lateral to the basal side, which led to the formation of structures resembling phagocytic cups all around the apoptotic cell (arrows), followed by the engulfment and apoptotic fragments (**Fig. 9C-E**). Occasionally, we observed formation contractile rings around apoptotic cell, which typically resulted in full or partial extrusion of apoptotic cells and were associated with the lack of full apical closure or failed corpse engulfment. **(Fig. 9F and Supplementary Fig. 10B-C)**.

To mechanistically assess the contribution of lamellipodia-based cell motility vs. actomyosin contractility in the elimination of dying cells by extrusion or by efferocytosis, we induced different types of cell death of single cells in co-cultures treated with the well-characterized Myosin II inhibitors blebbistatin and ML-7, the ROCK inhibitors Y-27632 and thiazovivin, the Rac1 inhibitor NSC-23766 and the CDC42 inhibitor ML-141 **(Fig. 9G)**. Consistently with previous work reporting cell extrusion, both myosin II (blebbistatin and ML-7) and ROCK inhibition (Y-27632 and thiazovivin) significantly delayed necrotic cell extrusion and increased the fraction of retained necrotic corpses 1-hour post cell death induction **(Fig. 9H and Supplementary Fig. 10D)**. On the other hand, blocking lamellipodia formation by Rac1 and CDC42 inhibitors by contrast only weakly affected necrotic cell extrusion. Blebbistatin and ML-7 treatment also affected apoptotic cells, by reducing the number of apoptotic cell fragments and moderately increased the fraction of fully or partially extruded apoptotic cells **(Fig. 9 I-J)**. Together, these data suggest that in Caco-2 cells, actomyosin contractility plays a dominant role in both necrotic cell extrusion and apoptotic cell fragmentation.

### Sphingosine-1-phosphate signaling is required for efferocytosis of apoptotic cells in monolayers

The release of sphingosines-1-phosphate from apoptotic cells and its sensing through S1P receptor 2 (S1PR2) by neighboring cells has been proposed to be a major driver of apoptotic cell extrusion^32–34^ **(Fig. 10A)**. We thus tested whether inhibition of this pathway blocks pyroptotic/necroptotic cell extrusion. Whereas treatment with JTE-013 (S1PR2 inhibitor) or SKI-II (Sphingosine kinase 2 inhibitor) slightly delayed extrusion of necroptotic cells (**Fig. 10 B-C**) from epithelia, it had no significant effect on the extrusion of the pyroptotic cells. This finding suggested that at least in Caco-2 cells S1P signaling was not the mayor pathway controlling extrusion of necrotic cells, and that despite the kinetic and mechanistic similarities, pyroptotic and necroptotic cells might produce different signals to drive their extrusion and elimination.

**Figure 10.**
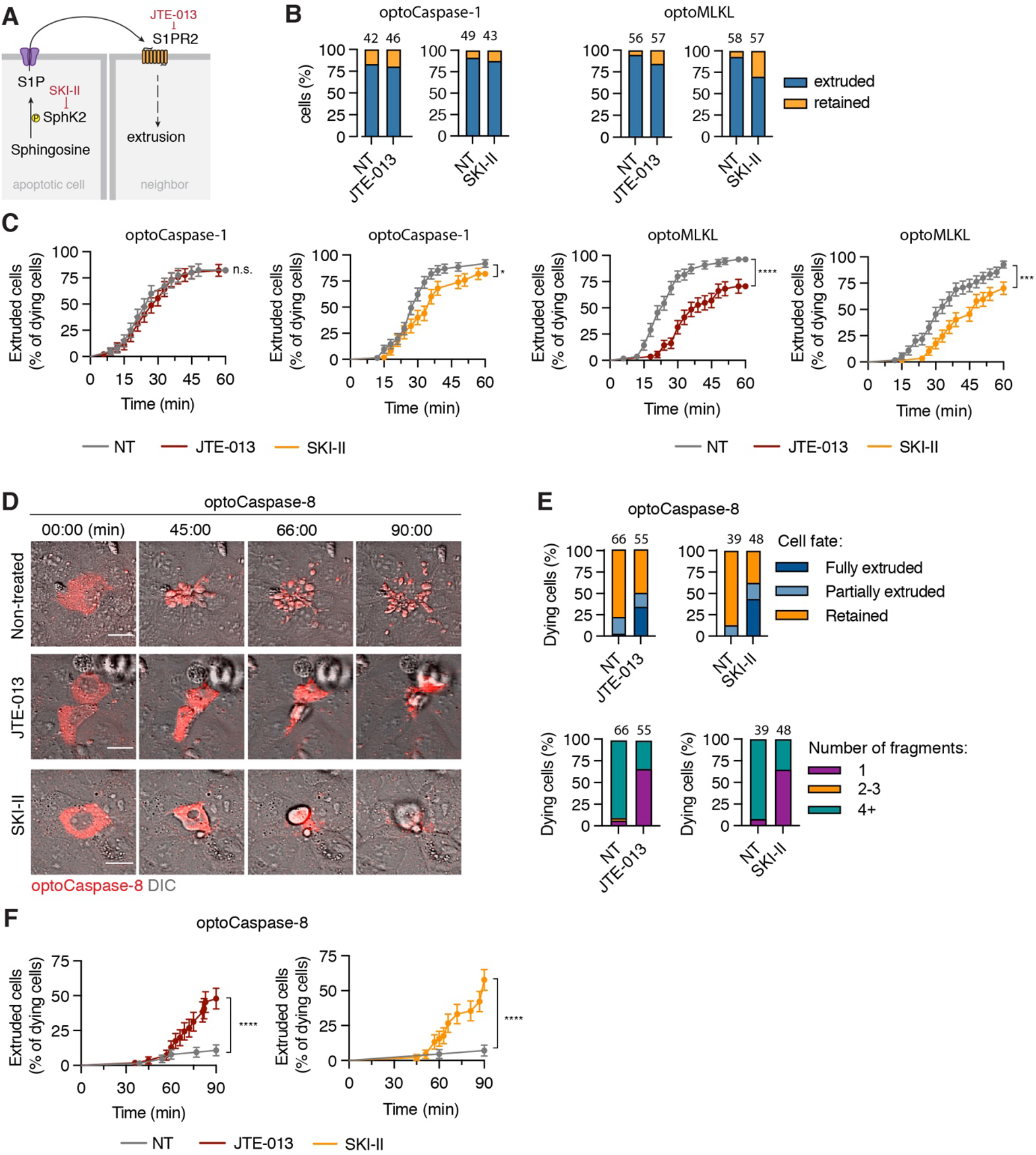
S1P signaling regulates apoptotic cell efferocytosis and necroptotic cell extrusion. **A,** Schematic illustration of S1P signaling. S1P is produced by Sphingosine phosphate kinase 2 (Sphk2) and released extracellularly, where it signals in an autocrine or paracrine manner by binding to Sphingosineig-1-phosphate receptor 2 (S1PR2). SKI-II and JTE-013 inhibit ShpK2 and S1PR2, respectively. **B,** Quantification of extruded vs. retained cells pyroptotic and necroptotic cells in the co-cultures treated with 30 *µ*m SKI-II or 20 *µ*m JTE-013 after 1h of light stimulation (7.8 mW/cm^2^ every 180 sec). **C,** Survival analysis extrusion probability of pyroptotic and necroptotic cells upon JTE-013 or SKI-II treatment. **D,** representative time-lapse images of optoCaspase-8-expressing Caco-2 cells undergoing light-induced apoptosis in presence of SKI-II or JTE-013. Scale bar: 20 *µ*m. **E-F,** Quantification and survival analysis of SKI-II or JTE-013-treated apoptotic cells in co-cultures. B, C, E and F are pooled from 3 independent experiments. C and F, * p<0.05, **** p<0.0001 (Mantel-Cox test).

By contrast, both JTE-013 and SKI-II treatment strongly affected the fate of apoptotic cells, as they resulted in reduced levels of apoptotic cell fragmentation (highlighted by the reduced numbers of apoptotic cell bodies being formed). This loss in cell fragmentation correlated with reduced uptake of apoptotic cells by neighbors and increase in number of extruded cells (**Fig. 10D-F and Supplementary movie 11**), suggesting that the inability of neighbors to fragment and engulf apoptotic cells or other apoptotic cell-derived signals may trigger their elimination by alternative mechanisms. Overall, these data suggest a previously unrecognized function for the S1P signaling pathway in apoptotic cell efferocytosis by epithelia, while also supporting a potential role of this pathway in necroptotic cell extrusion.

## Discussion

Here we report the development, optimization and characterization of a new toolset of <u>opto</u>genetically controlled <u>c</u>ell <u>d</u>eath <u>e</u>ffectors (optoCDEs) for the induction of three major programmed cell death modes – apoptosis, necroptosis and pyroptosis – in human and mouse cells and in zebrafish larvae. We show that optoCDEs allow the specific induction of these forms of cell death orthogonally to and with faster kinetics and higher efficiency than the endogenous cell death pathways, offering several advantages over other currently used methods of ‘clean’ cell death induction such as chemical activators or laser ablation. Compared to the forced dimerization and activation of caspases^9,35,36^ and RIP3/MLKL^37–40^ using modified FKBP domains, optoCDE not only excel in their rapid kinetics of activation and reversibility (inactivate within minutes of ceasing illumination), but above all in their use for single cell manipulation and imaging in 2D, 3D or *in vivo* settings. As we show, focused laser illumination yields superior spatiotemporal resolution that allows to rapidly and selectively kill individual cells without directly affecting the neighbors or the organism, even if these also express optoCDEs. Whereas other laser-based methods (such as laser ablation, laser-induced DNA-damage or photosensitizers, such as KillerRed) can also be used to induce cell death rapidly, in selected cells and *in vivo*^41–45^, they often fail to elicit specific cell death modes, rather relying on less specific types of cellular damage^46^. Laser ablation for example induces mixed apoptotic and necrotic phenotypes^7,8^, which limits its application to study the response of cells or tissue to specific types of PCD. By contrast, the optoCDE approach enables highly specific activation of selected cell death programs directly at effector protein level, increasing the specificity of cell death program and reducing the impact of endogenous regulation and inter-pathway crosstalk. While previous studies report several optogenetic strategies for apoptosis induction via clustering of death receptors^47^, recruitment of Bax^48^ to mitochondria or allosteric non-proteolytic activation of effector caspases via interdomain linker extension^49,50^, our system additionally benefits from utilizing similar molecular approach and illumination parameters for different effectors, enabling a more accurate comparison of signaling events induced by different types of cell death both *in vitro* and *in vivo*. Finally, the Cry2-based activation of cell death can be easily expanded to other cell death pathways that involve oligomerization of signaling components, such as lysosomal cell death or autophagy-driven cell death, and the recent identification of light-cleavable proteins allows to even extend this principle to cleavage-activated cell death executors, as recently shown for Bid and Gasdermin D^51,52^.

The development of optoCDEs offers new and unprecedented approaches to study cell death. Whereas “classical” methods for PCD induction often take hours and sometimes require additional priming or co-stimulation of the cells with multiple ligands, direct optogenetic activation of cell death on the effector level takes just few minutes, allowing for mechanistic studies of cells undergoing cell death with extremely high temporal resolution, and independently of confounding effects associated with regularly used activators (such as TNFa+SMAC or LPS+nigericin). Combined with the advanced live imaging techniques, this approach could offer unprecedented insights into the early cellular and metabolic events occurring during different types of PCD. The high reversibility of optoCDE on the other hand offer a new way to study cellular mechanisms useed to avoid or revert from cell death (anastasis)^53^, or the effects of sub-lytic/sub-lethal activation of cell death effectors like caspases and gasdermins. The mechanisms that drive and control sub-lethal cell death induction have recently gained attention, as it has been shown that low-level caspase activation results in ‘minority Mitochondrial outer membrane permeabilization’ (MiniMOMP) that drives DNA damage and genome instability^54^, or is important for differentiation^55^. Finally, the most valuable application, and the largest advantage over current methods, lies in the use of optoCDEs in the specific single cell death induction in 2D epithelia, 3D organoids and in live animals, and in the study of how these multicellular systems respond to distinct forms of PCD.

To specifically highlight this application of optoCDEs, we studied the differential fates of individual epithelial cells upon induction of apoptosis, necroptosis or pyroptosis. Dying cells release a number of ‘find-me’ and ‘eat-me’ signals that allows professional phagocytes to migrate towards and engulf dead cells (a process termed efferocytosis)^56^. Efferocytosis by non-professional phagocytes, including epithelial cells, has been reported as well^57^, but is less well understood as most epithelia, such as the gut epithelium, are thought to react to cell death by extruding dying cells^58^. For example, multiple in vitro studies clearly demonstrate extrusion of dying cells in response to different proapoptotic triggers, such as etoposide treatment^59,60^, UV illumination^28,32,61^ or starvation^62^. Our results using single cell killing by optoCDEs in human epithelial cell culture and in zebrafish skin however reveal that the type of cell death a dying cell undergoes might have a critical role in determining its fate. We find that necroptotic and pyroptotic cells are generally extruded from the epithelial layer, in line with previous study showing extrusion of epithelial cells after inflammasome activation^63,64^, and that this extrusion involves the simultaneous lamellipodia-based neighbor motility and contractile purse-string actin structures as reported previously for laser-ablated cells^22,30,31^. By contrast, the majority of apoptotic cells get fragmented and engulfed by their neighbors by a mechanism involving S1P signaling and formation of phagocytic cup-like structures. What determines these differential fates remains to be identified, but it is conceivable that necrotic and apoptotic cells release different signals/molecules or present different mechanical stimuli that elicit differential responses in their neighbors. While our findings differ strikingly from several studies that report extrusion of apoptotic cells^28,30–33,59,60,65,66^, these differences could be potentially due to different cell lines used, or the way cell death is induced.

In summary, optoCDEs provide a versatile and specific approach to investigate PCD pathways at a spatiotemporal level, not only in individual cells but also on their neighbors in a multicellular setting. This eliminates confounding effects from stimulatory signals, and thus might allow to identify and study new biological processes during PCD, and a better understanding of associated phenomena such as membrane repair, sub-lytic cell death, or cell extrusion and migration.

## Acknowledgements

This work was supported by a European Research Council Grant (ERC2017-CoG-770988-InflamCellDeath), Swiss National Science Foundation Project Grants (310030_175576 and 310030B_198005) and funding from the OPO Stiftung and Novartis to P.B. We thank Florence Morgenthaler and UNIL Cellular imaging facility (CIF), UNIL Flow Cytometry Facility, EMBL Advanced Light Microscopy Facility (ALMF) for technical assistance and to Darren Gilmour and Jonas Hartmann for providing the tg(6xUAS:mneonGreen-UtrCH) zebrafish line.

## Author contributions

K.S. and P.B conceptualized the study. K.S. and J.S. performed *in vitro* experiments; M.L and E.H. designed and performed *in vivo* experiments. All authors contributed to data analysis and manuscript writing.

## Materials and Methods

### Plasmids

The following constructs were obtained from Addgene: Cry2olig-mCherry (plasmid #60032), mouse RIP3-GFP (plasmid #41382) and Lifeact-miRFP703 (plasmid #79993) and used as the cloning templates. Vector containing human caspase-5 was obtained from Sino Biological (cat. #HG11152-M). The genes encoding human caspase-1, −4, −5, −8 or −9, RIP3 and MLKL, mouse caspase-1 and caspase-11 were amplified from human or mouse cDNA. Amplified fragments were fused C- or N-terminally to Cry2olig (as indicated in the text and figures) using overlap-extension PCR and subcloned into NheI/BstBI cloning sites of pLJM1 (Addgene plasmid 19319) for constitutive expression or EcoRI site of pLVX (TaKaRa, Cat. No. 632164) for doxycycline-inducible expression using InFusion HD cloning kit (TaKaRa). Additional GGGS linker was introduced between the Cry2olig and caspase/kinase domains or Cry2olig and mCherry to reduce potential sterical interference between domains. The exact construct design is indicated in the corresponding figures. For *in vivo* studies in zebrafish, the catalytic domain of zf caspa, caspb and caspase-8 was fused N-terminally to Cry2olig and subcloned into the pTH2 vector backbone containing a bidirectional heat-shock element (HSE) as promoter^26^, allowing heat-shock-induced expression of the cassette. The plasmids also contain the cmlc2:tagRFP as a transgenic marker and the insertion cassette is flanked by Tol2 sites for transgenesis^67^. All inserts were verified by sequencing to ensure the absence of unwanted mutations.

### Cell lines and tissue culture

HaCaT cell line was a generous gift of Gian-Paolo Dotto (University of Lausanne), Caco-2 cell line was a gift of Shaynoor Dramsi (Institut Pasteur, Paris), and MCF7 cell line was a gift of Nouria Hernandez (University of Lausanne). HT-29 cell line was purchased from Sigma (Cat. No. 91072201). Human GSDMD-transgenic cell line was described previously^21^. HEK293T, HeLa, Caco-2, HT-29, MCF7, 3T3 and RAW264.7 cells were maintained in Dulbecco’s Modified Eagle Medium (DMEM, Gibco) supplemented with the 10% fetal calf serum (FCS, Bioconcept), 200 U/ml penicillin and 200 μg/ml streptomycin (Bioconcept). HaCaT, U937 and THP-1 cells were maintained in RPMI medium supplemented with the 10% FCS and penicillin-streptomycin mix. Cells were maintained at 37°C, 5% CO_2_.

### Generation of stable cell lines

All stable cell lines were generated using an optimized lentiviral transduction protocol. For the production of lentiviral particles, 1 × 10^6^ HEK293T cells were transiently transfected using 1.9 μg of lentiviral expression vector (pLJM1 or pLVX), 1.9 μg of third generation packaging vector PsPax2, 0.2 μg of VSVg and 5 μl of JetPRIME transfection reagent. The supernatants were collected 24-48 h after transfection, filtered and either used immediately or preserved at −80 Co for the long-term storage. For transduction, approximately 1x 10^6^ cells of each cell line were spin-infected for 1 h at 1.9 × 10^3^ g (3000 rpm) in the presence of 10 μg/mL polybrene (Merck) to facilitate the infection. The virus-containing medium was replaced 24 h later, and cells were left to recover for additional 24-48 h in fresh medium. The stably transduced population of cells was selected using appropriate antibiotics (5 ug/ml puromycin or 50-100 μg/ml hygromycin Gold, both from Invivogen) for at least 5 days. To maintain stable expression, cell lines were subjected to the regular rounds of antibiotic treatment, and expression was monitored using fluorescence microscopy. Lifeact-GFP-expressing Caco-2 cells were additionally sorted based on GFP fluorescence to achieve a more uniform transgene expression level.

### Generation and lentiviral transduction of primary murine bone marrow derived macrophages

The primary bone marrow derived macrophages were isolated from 6-8-week-old wild-type C57BL/6 mice and differentiated in DMEM supplemented with 20% conditioned 3T3 cell supernatant (as a source of macrophage colony-stimulating factor M-CSF), 10% FCS, 10 mM Hepes HEPES (BioConcept), penicillin/streptomycin (Gibco) and non-essential aminoacids (Gibco). To express the optoCaspase-1/11, the macrophages were transduced with the lentiviral particles after 48 and 72 h of culture, as described previously^68^, and kept in the dark afterwards to avoid spontaneous construct activation. Photoactivation and imaging were performed on day 6-8.

### 3D cell culture

Single-cell Caco-2 suspension was mixed with 1.5-2 volumes of ice-cold Phenol Red Free Matrigel (Cornig), and a 40 μl drop of the resulting mixture was placed in the middle of each well of preheated 4-well μ-Slide (Ibidi), followed by immediate transfer at 37°C to induce Matrigel polymerization. 10 minutes later, 250-300 μl of complete medium was added to each well, and cells were maintained for 7-10 days to induce sphere formation, and the medium was replaced every 2-3 days. To induce the transgene expression, the medium was supplemented with 1 μg/ml doxycycline approximately for 16 h before the experiment. For visualization of pyroptotic cells, the imaging medium was supplemented with DRAQ7 for at least 1 h prior to imaging to allow for dye diffusion in Matrigel.

### Co-culture experiments

To generate mosaic co-cultures for cell extrusion studies, the optoCDE-expressing Caco-2 cells were trypsinized, mixed 1:20 with the wild-type Lifeact-GFP-expressing “neighbor” cells and plated on the tissue culture treated 8-well **µ**-chambers (ibidi) at a concentration of 2 × 10^5^ cells/well to form a confluent monolayer. OptoCDE expression was induced using 1 *µ*g/ml doxycyclin treatment overnight, and imaging/photostimulation were performed the following day. To accurately determine the dying cell and neighboring cell borders, the cell membranes were pre-stained with 5 ng/mL CellMask DeepRed (Invitrogen) for 60 min.

### Live cell imaging

All in vitro imaging experiments were performed using LSM800 point-scanning confocal microscope (Zeiss) equipped with 20× air immersion objective (for sphere imaging) and 63 × oil immersion Plan Apo objective (for all other experiments), 405, 488, 563 and 630 nm lasers. Temperature, CO_2_ and humidity were controlled throughout live imaging using an automated temperature control system and a gas mixer. For time-lapse imaging, Zeiss Definite Focus.2 system was used to ensure image stabilization. Images were acquired using Zen 2 software (Zeiss) at 16-bit depth, and acquisition settings were kept constant for the same type of experiments to aid statistical analysis.

### Photoactivation of optoCDE constructs

The photoactivation experiments were performed using 488 nm laser. The laser intensity was determined using FieldMaxII laser power meter (Coherent), and the light power density was calculated as follows ^69^: *Light power density* (W/cm^2^) = *Laser peak power [W] / effective focal spot area [cm^2^]^−1^*. The illumination settings were kept constant for the experiments of the same type, unless indicated otherwise.

For transient stimulation (Figures 3 and 4) cells were illuminated at selected timepoint with the indicated number of pulses using Zen “experimental regions” and “bleaching” functions. For sustained activation, the stimulation was performed repeatedly every 15 sec (unless otherwise indicated in figure legends) or every 90-180 sec (cell extrusion experiments). Initial testing and validation of the optoCDE constructs was performed in HEK293T cells. Briefly, 4 × 10^4^ cells were seeded in each well of collagen-coated 8-well *µ*-chamber (ibidi) and transfected with 150-300 ng of indicated construct using XtremeGene 9 transfection reagent (Roche) according to the manufacturer’s recommendation, and photostimulation/imaging was performed 24 h later. Prior to imaging, the cells were briefly washed, and the cell culture medium was replaced with the pre-warmed optiMEM. To visualize PS exposure and membrane permeabilization, the imaging medium was supplemented with CellTox Green (50’000 × dilution, Promega), 1 *µ*M DRAQ7 and/or 1 *µ*g/ PacificBlue-conjugated Annexin V (BioLegend). For cell death induction in transgenic cell lines, the cells were plated at a concentration of 1.5-2.0 × 10^5^ cells/well approximately 24 h before experiment, and optoCDE expression was induced by overnight treatment with 1 *µ*g/ml doxycycline. To avoid cell death due to spontaneous construct activation, cells were protected from light following transfection or expression induction, and all manipulations were performed under dim light.

### Activation and live imaging of optogenetic caspase variants in zebrafish

Selected larvae were heat-shocked at 2.5 dpf by incubation at 39°C for 30 min and kept under light shielding conditions. For imaging, larvae were anaesthetized by adding 40 *µ*g/ml ethyl m-aminobenzoate methanesulfonate (tricaine, Sigma) into the medium. They were then mounted in 1 % low–melting-point agarose (PEQ LAB Biotechnologie) on glass bottom dishes (MaTek). A Zeiss Biosystems 780 inverted confocal microscope was used for live imaging at RT. For skin and muscles cells we used a 40× water objective (LD C-Apochromat 40×/1.1 W Corr M27 or C-Apochromat 40×/1.2 W Corr M27; Zeiss Bio-systems). For whole larvae and overnight imaging, we acquired tile scans using a 20× air objective (Plan-Apochromat; Zeiss Biosystems) and stitching. The 488-laser was used at a power between 3-5 % to induce optogenetic activation and excitation of GFP. The 2-Photon laser was used at 950 nm wavelength and 20% laser intensity. For visualization of pyroptotic cells, the anaesthetizing medium and mounting agarose solution was supplemented with DRAQ7 (1:100).

### Phototoxicity and Cry2olig-induced cytotoxicity assessment

To ensure the absence of phototoxicity and Cry2olig-induced toxicity, the cells were transiently transfected with Cry2olig-mCherry plasmid, and whole imaging field was repeatedly scanned with the 488 nm laser of varying intensity (0.5 to 4.5% laser power, 4.8 to 40.9 mW/cm^2^) every 15 seconds for at least 1 h, and cell viability was assessed at end timepoint based on cell morphology and presence of Annexin V and CellTox staining. Based on these data, the illumination intensity range between 4.8 and 25 mW/cm^2^ was defined as non-toxic for the cells and used for all following experiments. To assess Cry2olig-induced toxicity, the percentage of mCherry-positive cells displaying altered morphology, Annexin V staining or CellTox influx was compared to the non-transfected (mCherry-negative) cells per same field of view.

### Generation and maintenance of Zebrafish strains

Experimental animals were cared about in accordance with EMBL guidelines and regulations and according to standard procedures ^70^. The following transgenic lines were created by co-injecting embryos at the one-cell stage with Opto-caspase plasmids with transposase mRNA (100 ng/*µ*l). Transgenic larvae were selected based on heart specific expression of tagRFP under the cardiac myosin light chain (cmlc2) promoter. The following lines were generated: *tg(mCherry-Cry2olig-caspa), tg(mCherry-Cry2olig-caspb)* and *tg(mCherry-Cry2olig-zfCaspase-8)*. Additionally we used the asc:asc-gfp^24^ for skin cell labeling. For visualizing cell boundaries the Opto-lines were crossed with *tg(Krt4:AKT-PH-GFP)*^71^. To follow actin dynamics in basal retaining cells we used the *tg(krt4:Gal4) line*^72^ crossed to tg(6xUAS:mneonGreen-UtrCH) generated in the lab of D.Gilmour by Jonas Hartmann. To inhibit pigmentation in larvae for imaging, we treated embryos with 0.2 mM 1-phenyl-2-thiourea (PTU; Sigma-Aldrich) in E3 medium.

### Drug treatment

The following inhibitors were used in the study: 20 μM Z-VAD-FMK (Invivogen), 1 μM GSK’872 (Selleck Chemicals), 5 μM Necrosulfonamide (Calbiochem), 50 *µ*M blebbistatin (Sigma), 30 *µ*M Y-27632 (StemCell), 15 *µ*M thiazovivin (StemCell), 50 *µ*M ML-141 (Sigma), 30 *µ*M ML-7 (Tocris), 200 *µ*M NSC-23766 (Tocris), 30 *µ*M JTE-013 (Sigma) and 30 *µ*M SKI-II (Tocris). All inhibitors were dissolved in DMSO and diluted to 10x concentration in OptiMEM, then added directly to the cells to achieve 1x concentration at least 1 h before imaging.

### Canonical and non-canonical inflammasome activation in U937 cells

For canonical inflammasome activation, PMA-differentiated U937 cells grown in 24-well plates were primed with LPS for 2 h, followed by the treated with the 15 μM Nigericin for 3 h to induce NLRP3 inflammasome activation. For non-canonical inflammasome activation, cells were primed with human IFNγ (10 ng/mL) overnight, and the LPS transfection was performed as described previously^3^. Briefly, 15 μg of smooth LPS from E. coli O111:B4 (Invivogen) was diluted in 200 μl of OptiMEM and incubated with Lipofectamine 2000 (5 μl per well) for 20 min at room temperature. Transfection complexes were then added directly on top of the cells in 200 μl of OptiMEM medium. To facilitate the LPS uptake, cells were centrifuged at 500 g for 15 minutes, followed by incubation at 37°C for 8 h.

### Optogenetic cell death induction in cell populations

The U937 cell were seeded onto black tissue culture treated 24-well plates (4titude) at concentration of 4 × 10^5^ cells/well. The differentiation towards the macrophage-like phenotype was induced using PMA treatment (5 ng/ml) for 24 h, after which the cells were washed once and left to recover for additional 24 h prior induction of expression. The HaCaT cells were seeded at a concentration of 4 × 10^5^ cells/well approximately 24 h before an experiment. Expression of the constructs was induced in both cell lines by treatment with the 1 μg/ml doxycycline overnight. To induce cell death, each of the wells was illuminated with the blue light LEDs (450 nm) using a custom-built Light Plate Apparatus device^2^. To manipulate light intensity and illumination duration, the custom illumination programs were created using Iris software (http://taborlab.github.io/Iris/).

### Western blotting

U937 were seeded at a density of 4 × 10^5^ cells/well and stimulated as indicated in the figure legends. After stimulation, the cells were lysed using pre-heated NuPage LDS sample buffer (Thermo Fisher Scientific) supplemented with 66 mM tris-Cl (pH 7.4), 2% SDS and 10 mM dithiothreitol (DTT). Cell supernatants were also collected, precipitated using methanol and chloroform and combined with cell lysates. Proteins were separated using 12% polyacrylamide gels and transferred onto a PVDF membrane using Trans-Blot Turbo (Bio-Rad). The following primary antibodies were used: rabbit anti-cleaved IL-1β (83186, CST; 1:1000), mouse anti-IL-1β (12242, CST; 1:1000), rabbit anti-GSDMD (ab210070, Abcam; 1:1000), rabbit anti-cleaved N-terminal GSDMD (ab215203, Abcam; 1:1000), mouse anti-caspase-1 (clone Bally-1 AG-20B-0048-C100, AdipoGen; 1:1000), mouse anti-mCherry (ab125096, Abcam; 1:2000), HRP-conjugated mouse anti-tubulin (ab40742, Abcam; 1:5000). Secondary antibodies conjugated to HRP (Southern Biotech; 1:10,000) were used for the chemiluminescent detection.

### Evaluation of cell lysis and IL-1β release

Cell lysis was assessed by measuring Lactate Dehydrogenase (LDH) activity in cell supernatants using the LDH cytotoxicity detection kit (TaKaRa). To obtain the positive control, the cells were fully lysed using 1% Tryton. To normalize for spontaneous cell lysis, the percentage of cell death was calculated as follows: (LDH_sample_ – LDH_negative control_)/(LDH_positive control_ – LDH_negative control_) × 100. The level of IL-1β in cell culture supernatants was measured using ELISA (R&D systems) according to the manufacturer’s protocol.

### Image analysis of *in vitro* data

All image analysis was performed using FiJi (https://imagej.net/Fiji) and Zen 2 (Zeiss) software. To aid visualization, the brightness and contrast of the representative images of the same panels were adjusted and set to similar values, and illumination correction for the DIC images was performed using Bandpass filter option (FiJi). For figures 1, 2, 4 and 6, the quantification of CellTox-positive and Annexin-positive cells at selected timepoints was performed manually due to the detachment and loss of late pyroptotic and necroptotic cells from imaging plane. Apoptotic cells were defined based on morphology (cell shrinking and blebbing) and Annexin V staining (Figure 1 and 4 and Supplemental figure 4). Quantification of DRAQ7-positive nuclei per field (Figure 3 B and C) was performed automatically using particle analysis tool and custom-written FiJi Macro, and the total number of DRAQ7+ nuclei per frame was divided by the total number of the mCherry-positive cells. The quantification of single-cell DRAQ7 intensities (Figure 3 and Figure 7) was performed by manually segmenting cell area and measuring the intensity density of selected regions over time, and intensity was normalized to the region intensity at t=0. Pyroptotic and necroptotic cell area was quantified by the manual segmentation and tracking of dying cell borders based on CellMask and mCherry signals and additionally validated using DIC channel. The extrusion time for Figures 8-10 was defined based on complete closure of the area previously occupied by the mCherry-positive cell, and for all experiments the t=0 was defined as the beginning of 488 laser stimulation. Pyroptotic and necroptotic cells were quantified as “extruded”, when the cell body and nucleus were fully removed from the plane of imaging, and remaining gap was completely closed by the neighbors (determined by membrane staining) and “retained” if the nucleus remained in the imaging plane at the end of experiment. Apoptotic cells were quantified as “fully extruded”, if the majority (over 80%) of apoptotic bodies were extruded by the end of experiment, “retained” if less then 20% of cell body was extruded, and “partially extruded”, if at least 20%, but not more than 80% of cell body remained within the monolayer.

### Quantification of apoptotic and pyroptotic events in vivo

Quantification of in vivo optoCaspase induced apoptotic and pyroptotic events in zebrafish keratinocytes was performed manually in several independent experiments for each of the three lines (*tg(mCherry-Cry2olig-caspa), tg(mCherry-Cry2olig-caspb)* and *tg(mCherry-Cry2olig-zf_caspase-8)*, and cells were classified as pyroptotic or apoptotic based on appearance of dead cells/cell debris. For the quantification of epithelial closure of either retention or extrusion of cells we counted the time starting at the frame before cell death related cell shape changes occurred to the timepoint at which surrounding cells have closed the gap completely at the apical site.

### Statistical analysis

All quantitative data analysis was performed using Microsoft Excel and GraphPad Prism 8. Statistical significances are referred as * (P <0.05), ** (P <0.01), *** (P <0.001) and **** (P <0.0001). For comparison of two groups, a two-tailed Student t-test was used. For comparison of three or more groups P-values were determined using the one-way analysis of variance (ANOVA) for multiple comparisons, and correction for the multiple comparisons was performed using Dunnett’s method. Comparison of extrusion probabilities of apoptotic, pyroptotic and necroptotic cells was performed using Mantel-Cox test.

