## Supplementary material for "Optogenetic activators of apoptosis, necroptosis and pyroptosis for probing cell death dynamics and bystander cell responses": Movie legends

**Description of additional supplementary files**

**Supplementary movie 1. Induction of pyroptosis by light-induced activation of optoCaspase-1, optoCaspase-4 or optoCaspase-5.** Time-lapse fluorescence confocal microscopy of GSDMD-transgenic HEK293T cells transiently transfected with indicated constructs and stimulated with blue light (5 mW/cm^2^) for 30 min. Merge of DIC, mCherry (marking optoCaspase or Cry2olig used as a control, red), CellTox (membrane permeabilization marker, green) and Annexin V (PS exposure marker, blue) is shown. Pyroptotic cells display characteristic morphological features, such as early membrane blebbing, swelling and ballooning, as well as nuclear rounding and condensation and acquisition of CellTox and Annexin staining. Cry2olig-expressing cells remain morphologically unchanged over the course of experiment. Scale bar, 20 µm.

**Supplementary movie 2. Sub-lytic induction of pyroptosis on a single-cell level.** Time-lapse fluorescence confocal microscopy showing HaCaT cells stably expressing optoCaspase-1 (red) that were transiently stimulated with the low dose of blue light a 3 min and left to recover for 1 h, after which a second round of 10 times more intensive light stimulation was performed. Inset (left) shows a cell displaying a sub-lytic membrane damage (as judged by the acquisition of low-intensity DRAQ7 staining, blue) after the first pulse. The cell transitions into “full” pyroptosis (gain of strong DRAQ7 signal, loss of cytosolic mCherry and membrane blebbing) after the 2^nd^ pulse of illumination. Scale bar, 20 µm.

**Supplemental movie 3. Optogenetic pyroptosis induction in single cells.** Time-lapse fluorescence confocal microscopy of HaCaT cells stably expressing optoCaspase-1 (red). Blue squares mark the regions which are transiently stimulated with the blue light at indicated timepoints. Pyroptotic cells rapidly gain DRAQ7 signal (blue) and are extruded from the monolayer by neighbors. Scale bar, 20 µm,

**Supplemental movie 4. Optogenetic pyroptosis induction in 3D organotypic cell cultures.** Time-lapse fluorescence confocal microscopy of Caco-2 cells grown in 3D cultures (spheres) in Matrigel and locally stimulated with the blue light at 5 min. Inset (left) shows a stimulated cell displaying membrane blebbing and swelling, gaining DRAQ7 signal (blue) and being extruded from the sphere wall into the lumen. Scale bar, 50 µm.

**Supplemental movie 5. Optogenetic induction of apoptosis via direct activation of optoCaspase-8 and optoCaspase-9.** Time-lapse fluorescence confocal microscopy of HEK293T cells transiently expressing optoCaspase-8 (red, left) and optoCaspase-9 (red, right) and stimulated with blue light. The appearance of early apoptotic features (cell shrinking and blebbing) precedes the PS exposure (Annexin V staining, blue). Scale bar, 20 µm.

**Supplemental movie 6. Optogenetic induction of apoptosis via direct activation of RIP3 and MLKL.** Time-lapse fluorescence confocal microscopy showing HEK293T cells expressing optoRIP3 (left, co-transfected with MLKL) or optoMLKL (right) and stimulated with blue light. Early necroptotic cells display only Annexin V staining (blue), while late necroptotic cells lose membrane integrity and gain CellTox signal (green). Scale bar, 20 µm.

**Supplemental movie 7. Optogenetic induction of pyroptosis in zebrafish larvae.** Time-lapse fluorescence confocal microscopy showing a zebrafish keratinocyte expressing opto-zfCaspa (red) and stimulated with blue light. Left, the brightfield channel. Pyroptotic cell can be seen extruded from the monolayer, followed by the gap closure. Scale bar, 20 µm.

**Supplemental movie 8. Optogenetic induction of apoptosis in zebrafish larvae.** Time-lapse fluorescence confocal microscopy showing a zebrafish keratinocyte expressing opto-zfCaspase-8 (red) and stimulated with blue light. Left, the brightfield channel. Apoptotic cell is fragmented into the apoptotic bodies, which are retained within the monolayer. Scale bar, 20 µm.

**Supplemental movie 9. Pyroptotic cell permeabilization in zebrafish.** Time-lapse fluorescence confocal microscopy showing a zebrafish keratinocyte expressing opto-zfCaspa (red) and stimulated with blue light. Left, the brightfield channel. Pyroptotic cell is rapidly extruded from the monolayer, which is followed by the rapid membrane permeabilization, loss of cellular content and gain of DRAQ7 signal (turquoise). Scale bar, 20 µm.

**Supplemental movie 10. Necrotic cell extrusion and apoptotic cell efferocytosis in human epithelial cells.** Time-lapse fluorescence confocal microscopy showing cells (red) undergoing optoCaspase-1-induced pyroptosis, optoMLKL-induced necroptosis or optoCaspase-8-induced apoptosis in presence of WT neighbors. Upper panels show CellMask (grey) and mCherry (red) signal, bottom panels show DIC and mCherry. Scale bar, 20 µm.

**Supplemental movie 11. Sphingosine-1-pathway regulates apoptotic cell efferocytosis by neighbors.** Time-lapse fluorescence confocal microscopy showing cells (red) undergoing optoCaspase-8-induced apoptosis in co-cultures which are either non-treated (left), treated with JTE-013 (S1PR2 inhibitor, middle) or SKI-II (Sphk2 inhibitor, right). Upper panels show CellMask (grey) and mCherry (red) signal, bottom panels show DIC and mCherry. Scale bar, 20 µm.
