## supplementary figures file for "Optogenetic activators of apoptosis, necroptosis and pyroptosis for probing cell death dynamics and bystander cell responses"

Supplementary material for

### Optogenetic tools for activation of programmed cell death

Shkarina *et. al.*

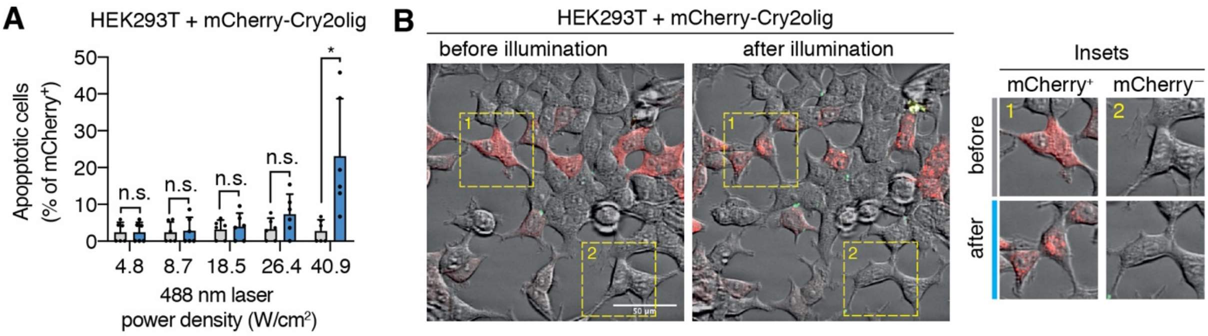


**Supplementary Figure 1. Blue light stimulation and Cry2olig activation do not induce cytotoxicity.**

**A,** To determine the phototoxicity threshold, HEK293T cells expressing mCherry-Cry2olig were illuminated with blue light of different power density every 15 sec for 1 h, and percentage of cells with altered morphology was quantified before and after illumination. **B**, Representative images of cells from A before and after 1 hour after illumination with blue light (5 mW/cm$^2$). Insets show the morphology of transfected (mCherry$^+$) and non-transfected (mCherry$^-$) cells.

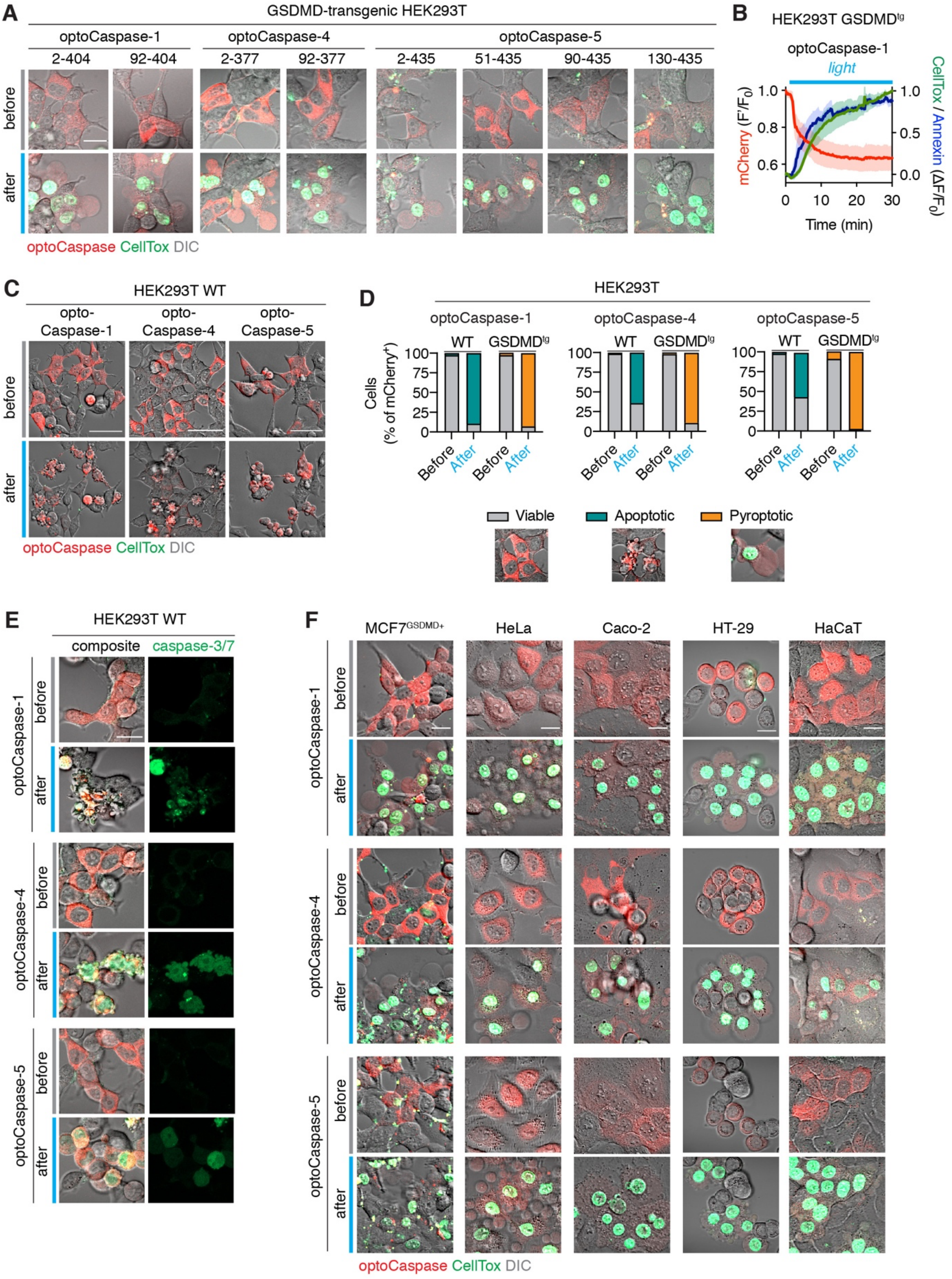


**Supplementary Figure 2. Validation of inflammatory optoCaspases.**

**A,** Representative images of HEK293T cells expressing different optoCaspase-1, -4 and -5 versions before and after illumination (488 nm, 5 mW/cm$^2$). **B,** Normalized fluorescence intensity of the mCherry, Annexin-V and CellTox signals in cells expressing CARD-deficient optoCaspase-1 and undergoing light-induced pyroptosis. Mean ± s.e.m., n = 5 cells. **C,** Representative images of wild-type (i.e. naturally GSDMD-deficient) HEK293T cells expressing optoCaspase-1, -4 or -5 (red) before and after illumination (488 nm, 5 mW/cm$^2$). **D**, Quantification of apoptotic, pyroptotic or viable cells before and after illumination among wild-type or GSDMD$^{tg}$ HEK293T cells expressing optoCaspase-1, -4 or -5. Images show morphological features and CellTox staining of the 3 categories of cells (viable, apoptotic and pyroptotic). **E,** Representative images of wild-type HEK293T cells expressing optoCaspase-1/-4/-5 and the genetically encoded caspase-3/7 activity reporter VC3AI (green) before and after illumination (488 nm, 5 mW/cm$^2$). **F,** Representative images of different human epithelial cell lines expressing optoCaspase-1/-4/-5 before and after 1 h of illumination (488nm, 5 mW/cm$^2$). Scale bars, 20 $\mu$m. All data are representative of at least 3 independent experiments.

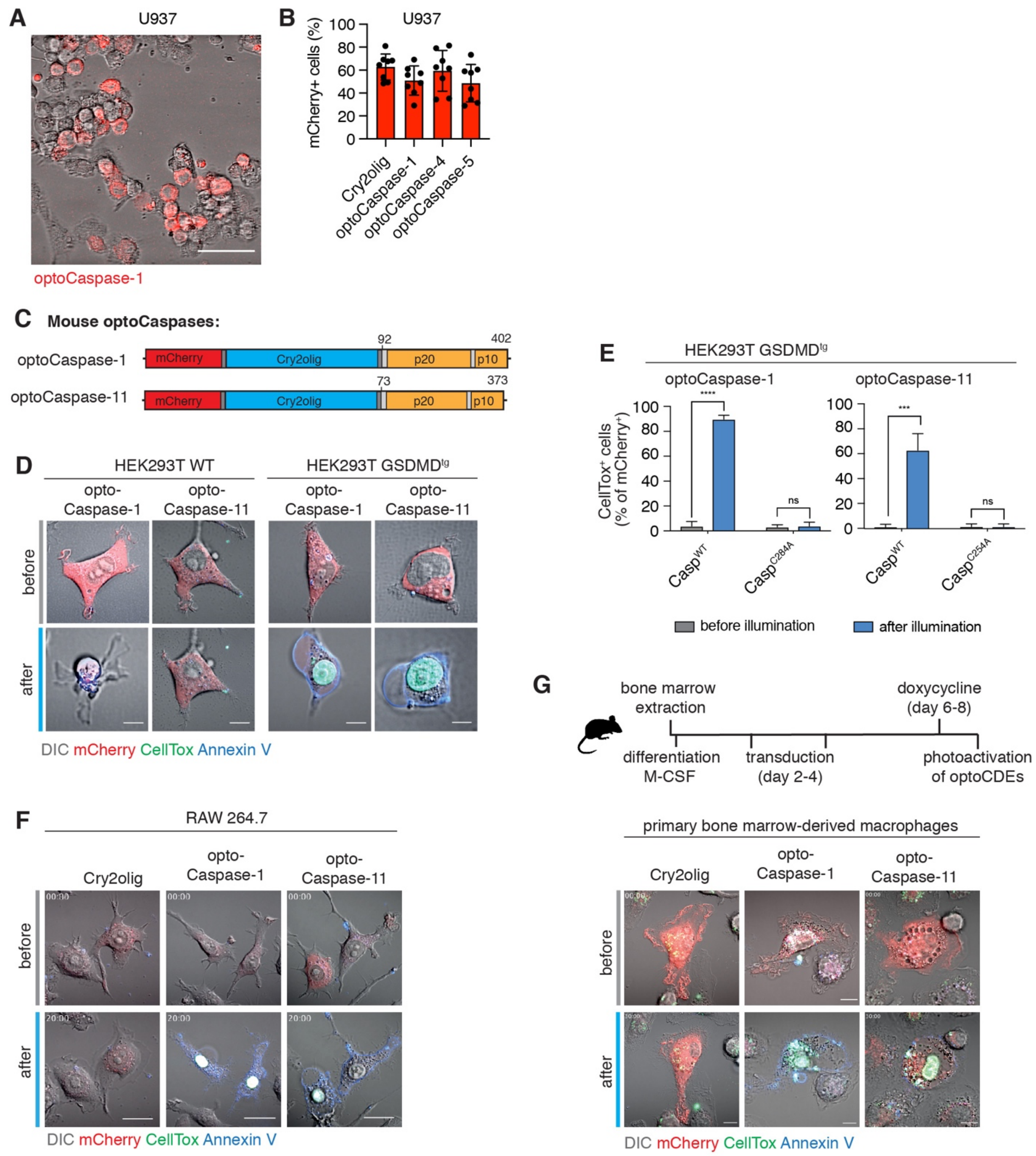


**Supplementary Figure 3. Optogenetic induction of pyroptosis in human and mouse macrophage-like cells.**

**A,** Representative image showing heterogenous expression levels in U937 cells stably transduced with optoCaspase-1 after doxycycline (dox) treatment. Scale bar, 50 $\mu$m. **B,** Percentage of mCherry-positive cells in differentiated dox-treated U937 cell lines expressing indicated constructs. Data is pooled from three independent experiments, 4 fields of view per experiment, mean ± s.d. **C,** Schematic presentation of mouse optoCaspase-1 and optoCaspase-11 constructs. Both constructs contain CARD-deficient mouse caspase-1 or 11 proteins N-terminally fused to mCherry-tagged Cry2olig via a GGGS linker. **D,** Representative images of wild-type or GSDMD-transgenic HEK293T cells expressing mouse optoCaspase-1 or optoCaspase-11 before or 30 min after illumination (488 nm, 5 mW/cm$^2$ every 15 sec). Cells were imaged in presence of CellTox (green) and Annexin-V (blue) to visualize membrane permeabilization and PS exposure. Scale bars, 10 $\mu$m. **E,** Percentage of CellTox-positive GSDMD$^{tg}$ HEK293T cells expressing wild-type or catalytically-deficient optoCaspase-1 or optoCaspase-11 before and after illumination (488 nm, 5 mW/cm$^2$ every 15 sec). Mean ± s.d., pooled from 3 independent experiments. **** $p < 0.0001$, *** $p < 0.001$, n.s. – non-significant (two-tailed t-test). **F-G,** Representative images of RAW 264.7 cells and primary murine bone marrow derived macrophages (BMDMs) expressing Cry2olig, optoCaspase-1 or optoCaspase-11 before and after illumination (488 nm, 5 mW/cm$^2$) in presence of CellTox and Annexin V. Scale bars, 20 $\mu$m (F) and 10 $\mu$m (G).

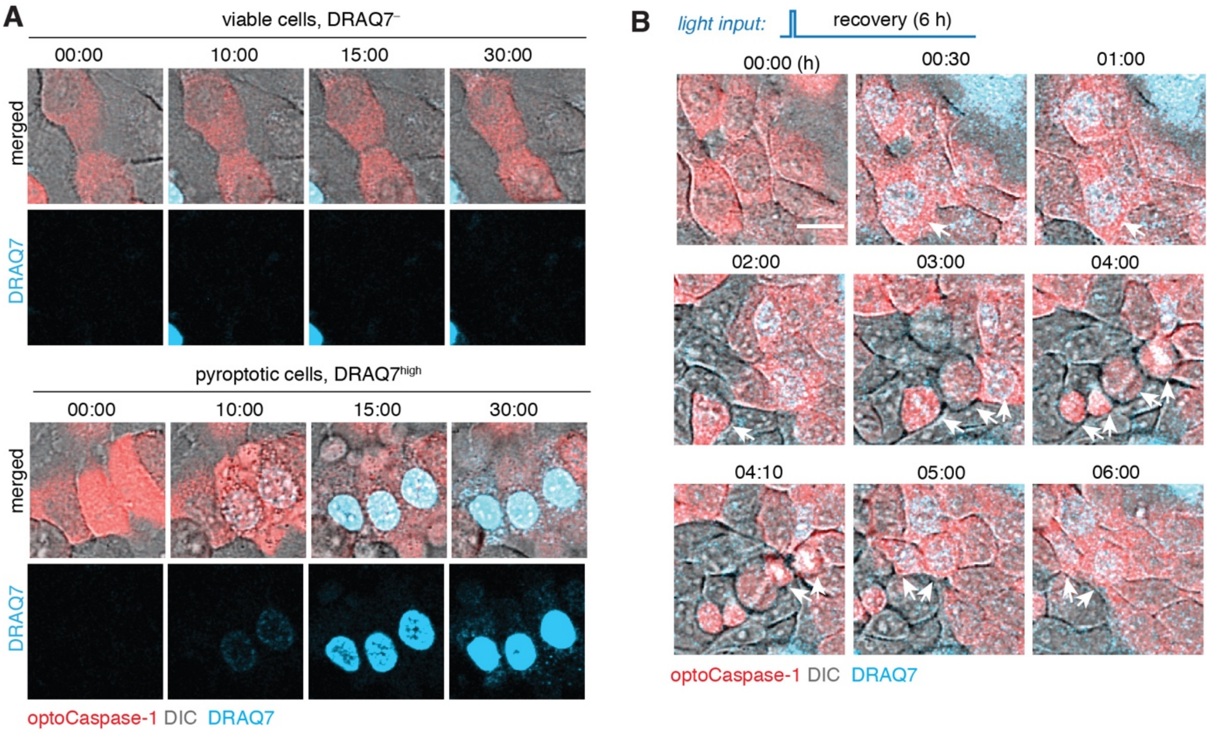


**Supplementary Figure 4. Sub-lytic induction of pyroptosis.**

**A,** Representative images of cells not affected by the first illumination pulse (viable, DRAQ7$^{-}$) or undergoing complete pyroptosis (pyroptotic, DRAQ7$^{high}$). Related to Figure 3E. **B,** HaCaT cells expressing optoCaspase-1 were transiently illuminated with low-intensity 488 nm light (3x0.2 mW/cm$^2$) at t=3min and imaged for 6 h. DRAQ7 influx (turquoise) marks the cells with initial sub-lytic membrane permeabilization, which was then repaired. Arrows indicate dividing parent cell and its daughters. Scale bar, 20 $\mu$m. Representative of 2 independent experiments.

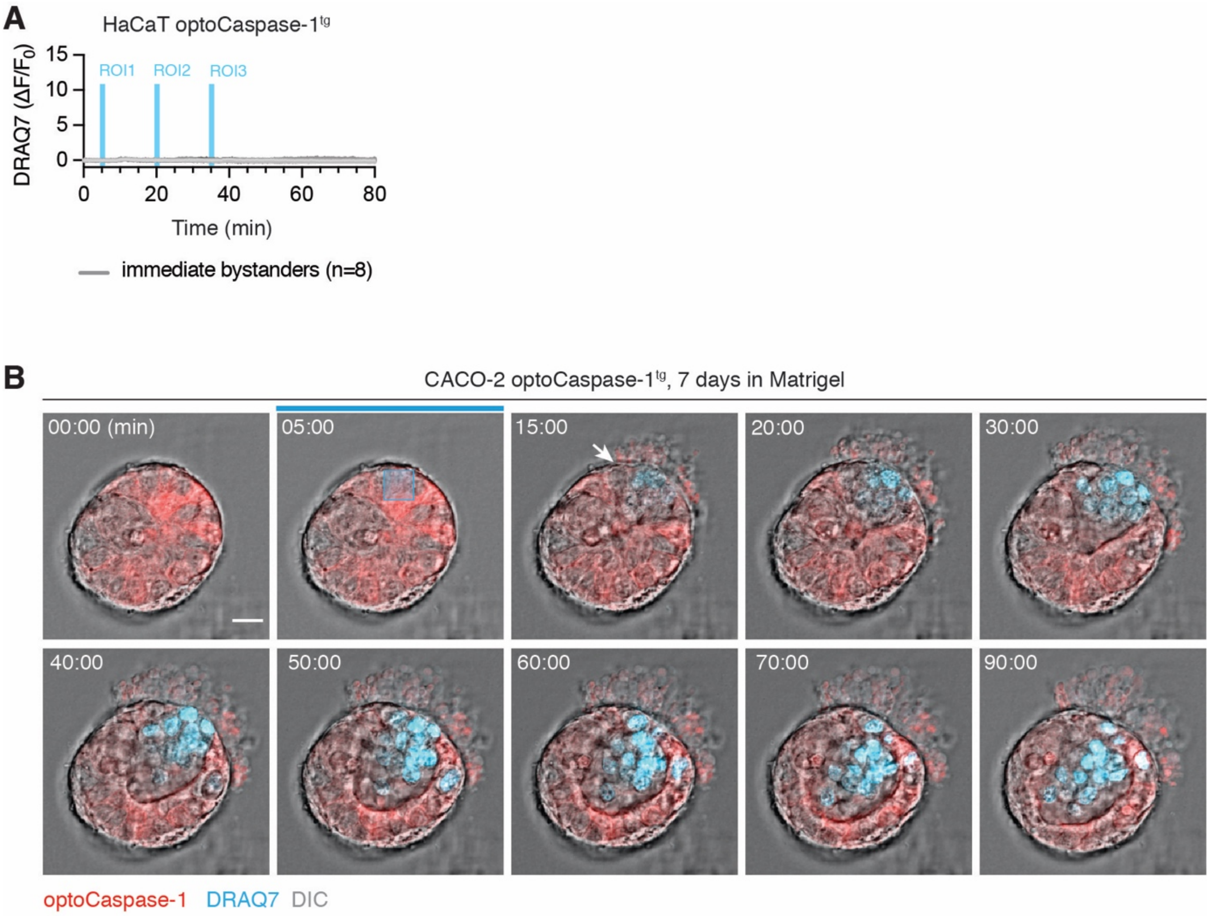


**Supplementary Figure 5. Single-cell ablation in 2D and 3D cell cultures.**

**A,** Normalized nuclear DRAQ7 intensity of 8 unstimulated directly adjacent neighbors of the cells selectively stimulated with blue light. Related to Figure 4A. Grey lines indicate individual cell traces. Vertical blue lines indicate the timing of each of the three local blue light stimulations. **B,** Representative time-lapse images of partial cell ablation in non-lumenized Caco-2 spheroid. Blue sphere indicates the region where the stimulation was performed at t=5 min. Scale bar, 50 *μ*m. Representative of 3 independent experiments.

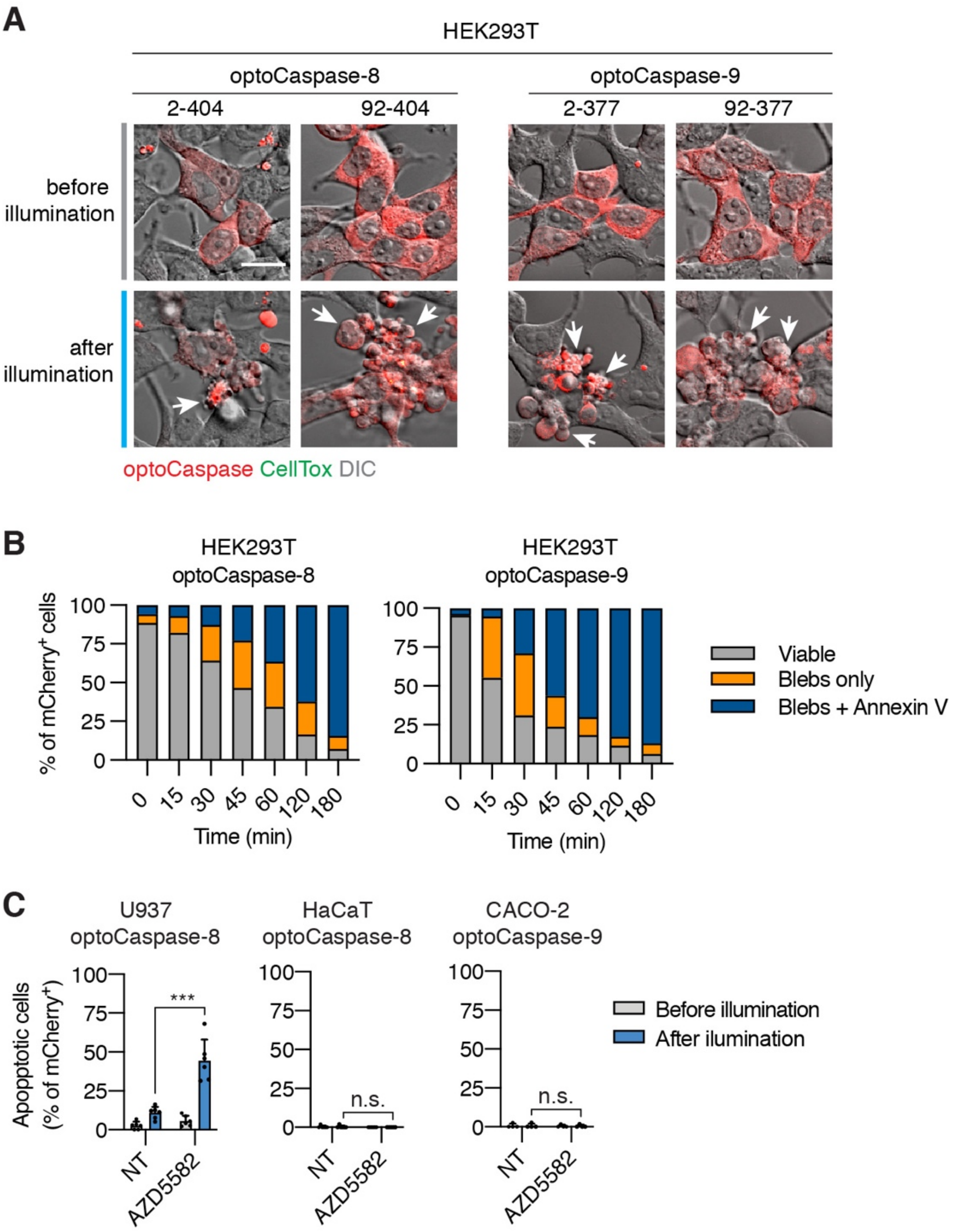


**Supplementary Figure 6. Optogenetic induction of apoptosis.**

**A,** Representative images of HEK293T cells expressing full-length or CARD/DED-deficient optoCaspase-8 or optoCaspase-9 before and after illumination (488 nm, 5 mW/cm$^2$ every 15 sec). Arrows indicate apoptotic blebbing. **B,** Quantification of cells from A expressing optoCaspase-8 or optoCaspase-9 and displaying apoptotic blebbing alone, or blebbing and Annexin V staining. Cells were stimulated with blue light every 3 min sec, and images were acquired for 3 h. **C,** U937 or HaCaT cells expressing optoCaspase-8, or Caco-2 cells expressing optoCaspase-9 were treated with the SMAC mimetic AZD5582 and stimulated with blue light for 1 h, and percentage of cells displaying apoptotic morphology was quantified before and after illumination. Mean ± s.d., pooled from 3 independent experiments. *** $p < 0.001$, n.s. – non-significant (two-tailed t-test).

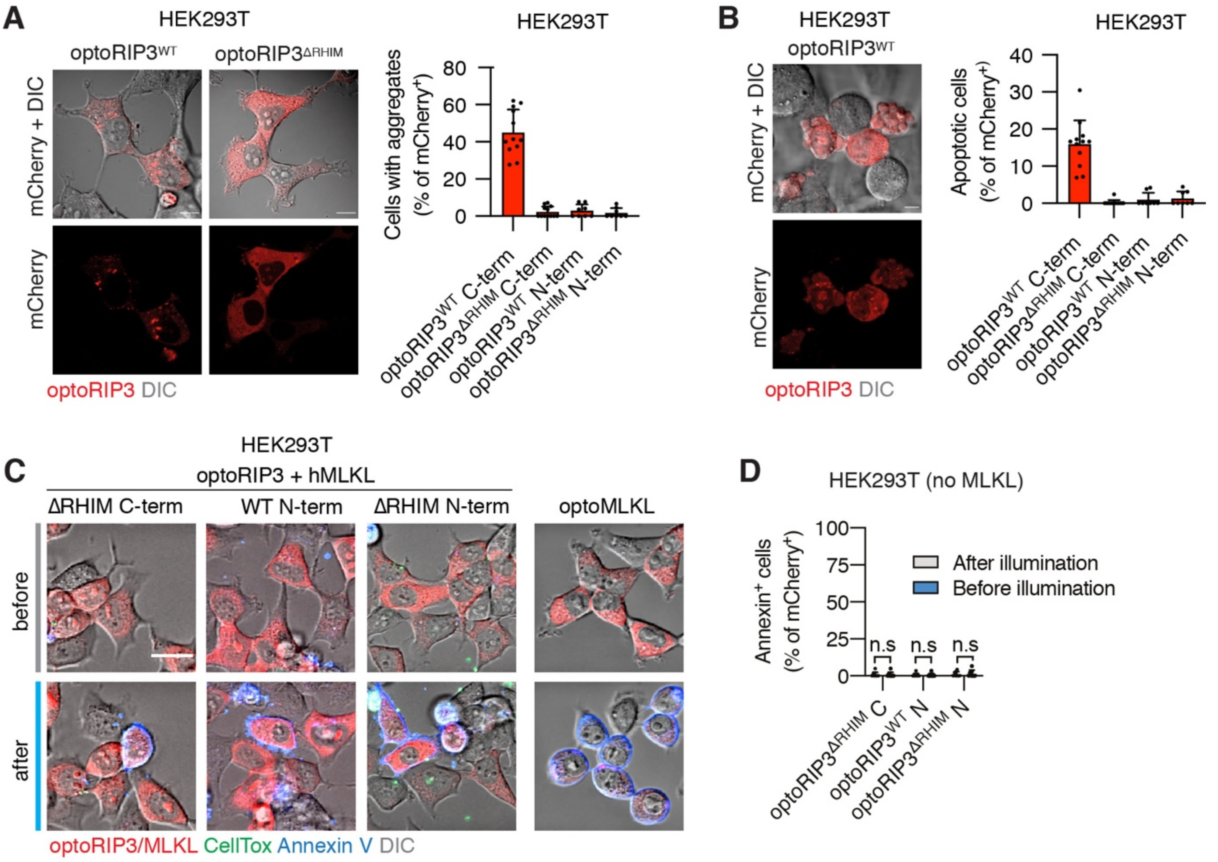


**Supplementary Figure 7. Optogenetic activation of necroptosis.**

**A,** Representative images of HEK293T cells expressing wild-type or RHIM-deficient optoRIP3 (C-terminally fused to Cry2olig) in absence of blue light illumination (left). Percentage of cells with mCherry clusters in HEK293T cells expressing different optoRIP3 versions (right). **B,** Representative images of cells expressing wild-type C-terminal optoRIP3 and displaying apoptotic morphology in absence of blue light (left). Percentage of spontaneously apoptotic cells in HEK293T cells transfected with different optoRIP3 versions (right). **C,** Representative images of HEK293T cells expressing different optoRIP3 versions (co-transfected with human MLKL) or optoMLKL before and after stimulation with blue light (5 mW/cm$^2$). Cells were imaged in presence of Annexin V (blue) and CellTox (green) to visualize early (pre-lytic) and late (lytic) necroptosis. **D,** Percentage of Annexin-V-positive HEK293T cells expressing optoRIP3 versions before and after illumination, in absence of MLKL co-expression. Data are pooled from 3 independent experiments, mean ± s.d., n.s. – non-significant (two-tailed t-test). Scale bars, 10 (A and B) or 20 (C) $\mu$m.

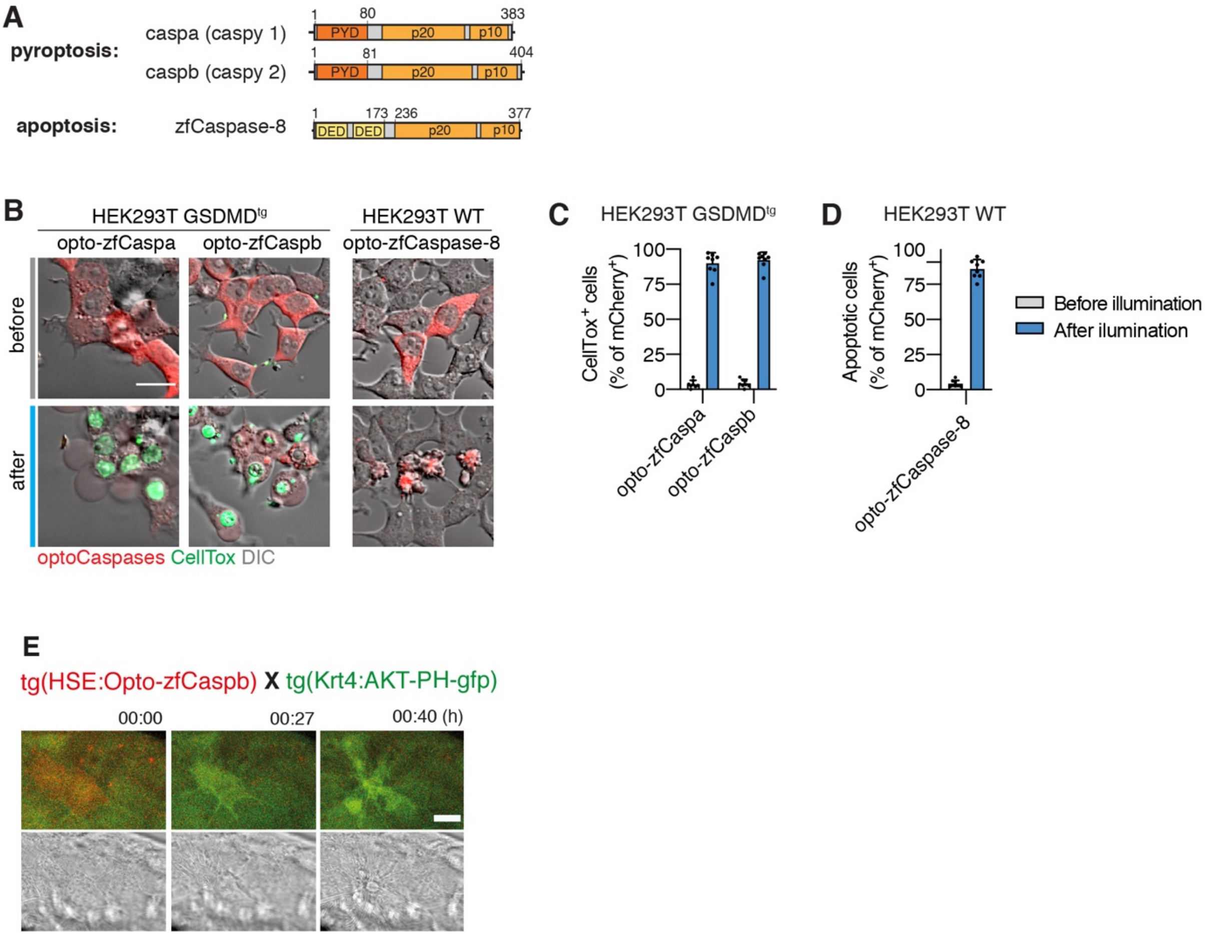


**Supplementary Figure 8. Activation of zebrafish caspases by optogenetics**

**A,** Domain organization of zebrafish inflammatory (caspa and caspb) or apoptotic (caspase-8) caspases. **B,** Representative images of human GSDMD$^{tg}$ HEK293T cells expressing opto-zfCaspa, opto-zfCaspb or opto-zfCaspase-8 before and after 30 min of blue light illumination (5 mW/cm$^2$). CellTox staining (green) indicates membrane permeabilization. Scale bar, 20 $\mu$m. **C-D,** Percentage of pyroptotic (C) or apoptotic (D) opto-zfCaspa or opto-zfCaspb-expressing (C) or opto-zfCaspase-8-expressing (D) GSDMD$^{tg}$ HEK293T cells before and after blue light illumination (5 mW/cm$^2$). Data is pooled from 3 independent experiments, mean ± s.d. **E,** Representative time-lapse images of a keratinocyte, after activation of opto-zfCaspb. Scale bar, 20 $\mu$m.

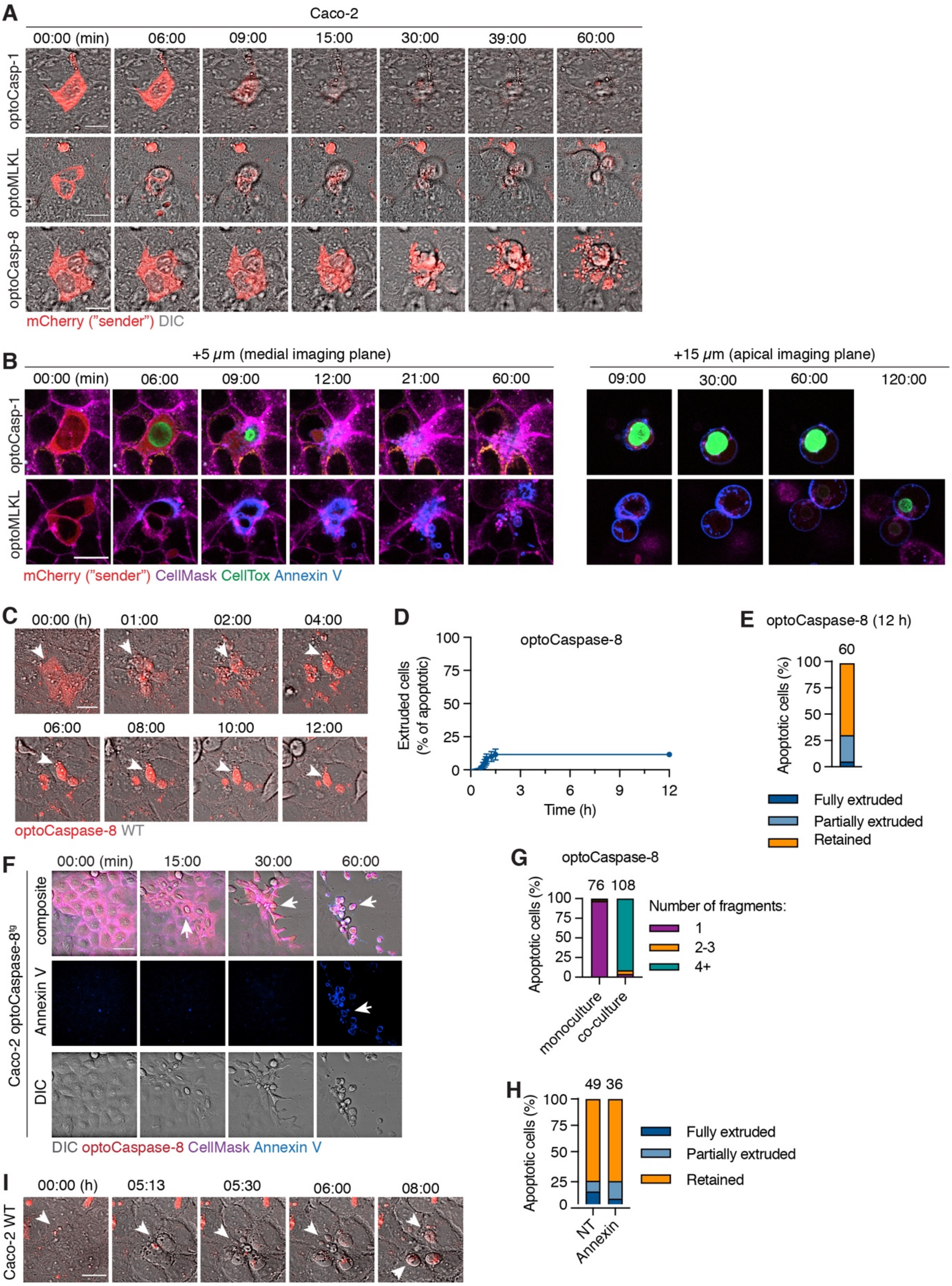


**Supplementary Figure 9. Differential response to apoptotic and necrotic cell death in epithelial cell populations.**

**A,** Representative time-lapse images (showing DIC and mCherry channels) of Caco-2 cells undergoing light-induced pyroptosis, apoptosis and necroptosis in co-cultures with wild-type cells (related to Fig. 8A-B). **B**, Representative time-lapse images showing CellMask (purple), Annexin-V (blue) and CellTox (green) staining in pyroptotic (optoCaspase-1) and necroptotic (optoMLKL) cells being extruded from monolayer of WT cells. Left, images of the cells at the medial plane (5 $\mu$m above basal surface), right, apical plane (15 $\mu$m above basal surface, showing extruded cells). For optoMLKL cells, imaging was performed for 120 min to show CellTox influx during lytic necroptotic stage. **C,** Time-lapse images of apoptotic cell (induced by optoCaspase-8 activation) in co-culture. Imaging was performed every 3 min for 12 h in total. Arrows indicate persisting apoptotic bodies in neighbors. **D-E,** Percentage of extruded apoptotic cells during 12 h post apoptosis induction. (D) Cells were classified as "extruded", if more than 50% of cell body or apoptotic fragments were expulsed from the monolayer. (E) full and partial extrusion were defined as in Fig. 8. **F,** Representative time-lapse images of optoCaspase-8-transgenic CACO-2 cells undergoing apoptosis in absence of neighboring cells. Annexin V staining (blue) marks PS exposure during late timepoint. Arrows indicate characteristic apoptotic features: initial cell contraction (15 min), cell shrinking and detachment (30 min), rounding up and PS exposure (60 min). **G,** Percentage of fragmented apoptotic cells in optoCaspase-8-transgenic Caco-2 cells cultured either alone (monoculture) or together with WT Caco-2 and stimulated with blue light for 1 h. **H,** Percentage of extruded, partially extruded apoptotic cells in co-cultures treated with Annexin-V (10 $\mu$g/mL). I, representative time-lapse images of spontaneous apoptosis and apoptotic corpse efferocytosis in wild-type Caco-2. Scale bars, 20 $\mu$m. Data are pooled from 2 (D-E and H) or 3 (G) independent experiments. All images are representative of at least 3 independent experiments.

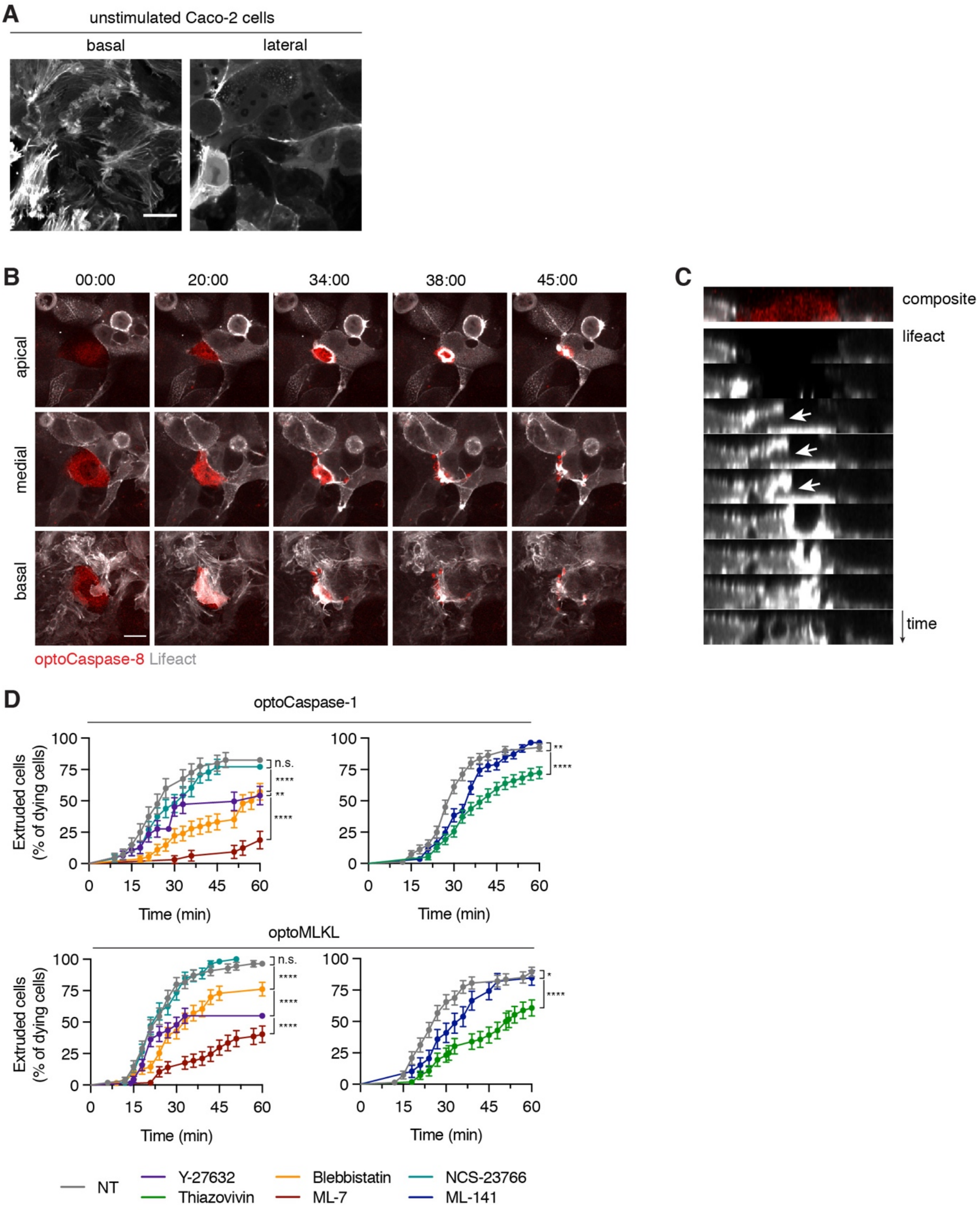


**Supplementary Figure 10. Apoptotic and necrotic cell death trigger differential cytoskeletal rearrangement.**

**A,** Representative images of lifeact-GFP distribution in non-stimulated CACO-2 cells. Basal plane (0 $\mu$m) shows randomly oriented lamellipodia, and medial plane (+5 $\mu$m) shows cortical actin enrichment at the cell junctions). **B,** Representative time-lapse images of actin rearrangement in neighboring cells during apoptotic cell extrusion. Scale bar, 20 $\mu$m. Cells were stimulated and images were acquired as in Fig. 8 A-C. **C,** Reconstructed side view (xz) of time-lapse series from B. Arrows indicate incomplete phagocytic cup formation during failed corpse engulfment. **D**, Percentage of extruded cells over time. Cell death was induced by illumination as described previously in the co-cultures treated with the indicated inhibitors. The time of extrusion was determined by complete gap closure in the imaging plane, determined by the CellMask signal. All data are pooled from 3 independent experiments, mean ± s.e.m.,
